# scMET: Bayesian modelling of DNA methylation heterogeneity at single-cell resolution

**DOI:** 10.1101/2020.07.10.196816

**Authors:** Chantriolnt-Andreas Kapourani, Ricard Argelaguet, Guido Sanguinetti, Catalina A. Vallejos

## Abstract

High throughput measurements of DNA methylomes at single-cell resolution are a promising resource to quantify the heterogeneity of DNA methylation and uncover its role in gene regulation. However, limitations of the technology result in sparse CpG coverage, effectively posing challenges to robustly quantify genuine DNA methylation heterogeneity. Here we tackle these issues by introducing scMET, a hierarchical Bayesian model which overcomes data sparsity by sharing information across cells and genomic features, resulting in a robust and biologically interpretable quantification of variability. scMET can be used to both identify highly variable features that drive epigenetic heterogeneity and perform differential methylation and differential variability analysis between pre-specified groups of cells. We demonstrate scMET’s effectiveness on some recent large scale single cell methylation datasets, showing that the scMET feature selection approach facilitates the characterisation of epigenetically distinct cell populations. Moreover, we illustrate how scMET variability estimates enable the formulation of novel biological hypotheses on the epigenetic regulation of gene expression in early development. An R package implementation of scMET is publicly available at https://github.com/andreaskapou/scMET.

## Introduction

DNA methylation (DNAm) at cytosine residues plays an important role in the regulation of gene expression (Jaenisch and Bird, 2003). It is also critical for a broad range of biological processes, including X-chromosome inactivation, genomic imprinting and cancer (Avner and Heard, 2001; Baylin and Jones, 2011; Reik and Walter, 2001). The gold standard approach to profile DNAm at single-base resolution is to treat DNA with sodium bisulphite, which efficiently converts unmethylated cytosines to uracils, while leaving methylated cytosines unmodified (Krueger *et al.*, 2012). Although bulk bisulphite sequencing (BS-seq) experiments have paved the way for mapping the methylome landscape across different tissues, they fall short of explaining the inter-cellular methylation heterogeneity and quantifying its dynamics in a variety of biological contexts (Schwartzman and Tanay, 2015).

More recently, advances in sequencing technologies have enabled the development of protocols that profile DNAm with single-cell resolution (e.g. scBS-seq, Guo *et al.*, 2013; Smallwood *et al.*, 2014) and multiplexing protocols offer scalability to thousands of cells in a single experiment (Luo *et al.*, 2017; Mulqueen *et al.*, 2018). In contrast to gene expression signatures from scRNA-seq experiments, which are influenced by the environment, DNA methylation profiles are highly distinct between cell types and stable across individuals and over the life span (Lister *et al.*, 2013; Mo *et al.*, 2015). Moreover, whilst scRNA-seq assays might fail to capture information about genes with moderate expression levels, cell-level measurements of DNAm offer a more complete coverage across genomic regions (Luo *et al.*, 2017). However, due to the small amounts of initial genomic DNA and the destructive nature of bisulphite on nucleic acids, the output data are often noisy and extremely sparse; that is, a large proportion of CpG dinucleotides is not observed (ranging from 80% to 95%). While tailored computational imputation methods such as Melissa (Kapourani and Sanguinetti, 2019) and DeepCpG (Angermueller *et al.*, 2017) might ameliorate the sparsity problem, disentangling genuine epigenetic variability from technical biases remains a formidable problem.

Here we present scMET, a Bayesian framework that addresses the statistical challenges associated with sparse scBS-seq data and provides novel functionality that is tailored to single-cell level datasets. To overcome sparsity, scMET aggregates the input data within regions (hereafter also referred to as genomic *features*): either by combining CpG information in a sliding window approach or using pre-annotated contexts, such as promoter regions or enhancers (Gravina *et al.*, 2016; Smallwood *et al.*, 2014). To dissect genuine epigenetic variability from the many confounding technical biases, scMET adopts a hierarchical model specification which shares information across cells and genomic features, whilst incorporating feature-level characteristics (e.g. CpG density). Critically, scMET introduces residual *overdispersion* estimates as a measure of DNAm variability that is not confounded by differences in mean methylation. These estimates can be used to perform differential DNAm variability testing among groups of cells, embracing the cellular resolution of the data to provide novel insights which are not possible using traditional differential mean tests on bulk data (Lähnemann *et al.*, 2020). scMET can also identify highly variable features (HVFs) which, among others, can be used as input for unsupervised clustering analyses.

scMET scales readily to thousands of cells and features, making it a powerful tool for large scale single-cell epigenetic studies. Our results both on simulated and real datasets demonstrate that it can accurately and robustly quantify DNAm heterogeneity. Results on two recent large scale datasets show that scMET detects biologically relevant highly variable features which result in improved clustering performance. In addition, we show that scMET can facilitate the interrogation of single-cell multi-omics assays, yielding novel biological hypotheses on the role of epigenetic variability in gene regulation in early development.

## Results

### Quantifying cell-to-cell DNAm heterogeneity with scMET

To disentangle technical from biological variability and overcome data sparsity, scMET couples a hierarchical beta-binomial (BB) model with a generalised linear model (GLM) framework (Fig. 1a-b). For each cell *i* and feature *j*, the input for scMET is the number of CpG sites that are observed to be methylated (*Y_ij_*) and the total number of sites for which methylation status was recorded (*n_ij_*). The BB model uses feature-specific mean parameters *μ_j_* to quantify overall DNAm across all cells and biological *overdispersion* parameters *γ_j_* as a proxy for cell-to-cell DNAm heterogeneity. The latter capture the amount of variability that is not explained by binomial sampling noise, which would only account for technical variation. Hence, *γ_j_* is akin to the overdispersion term used in negative binomial models for RNA-seq data (e.g. Love *et al.*, 2014). Although BB models have been developed for bulk DNAm data (e.g. Dolzhenko and Smith, 2014; Feng *et al.*, 2014), they typically use data from individual CpG sites as input; a strategy prone to fail for the highly sparse scBS-seq data.

**Figure 1:**
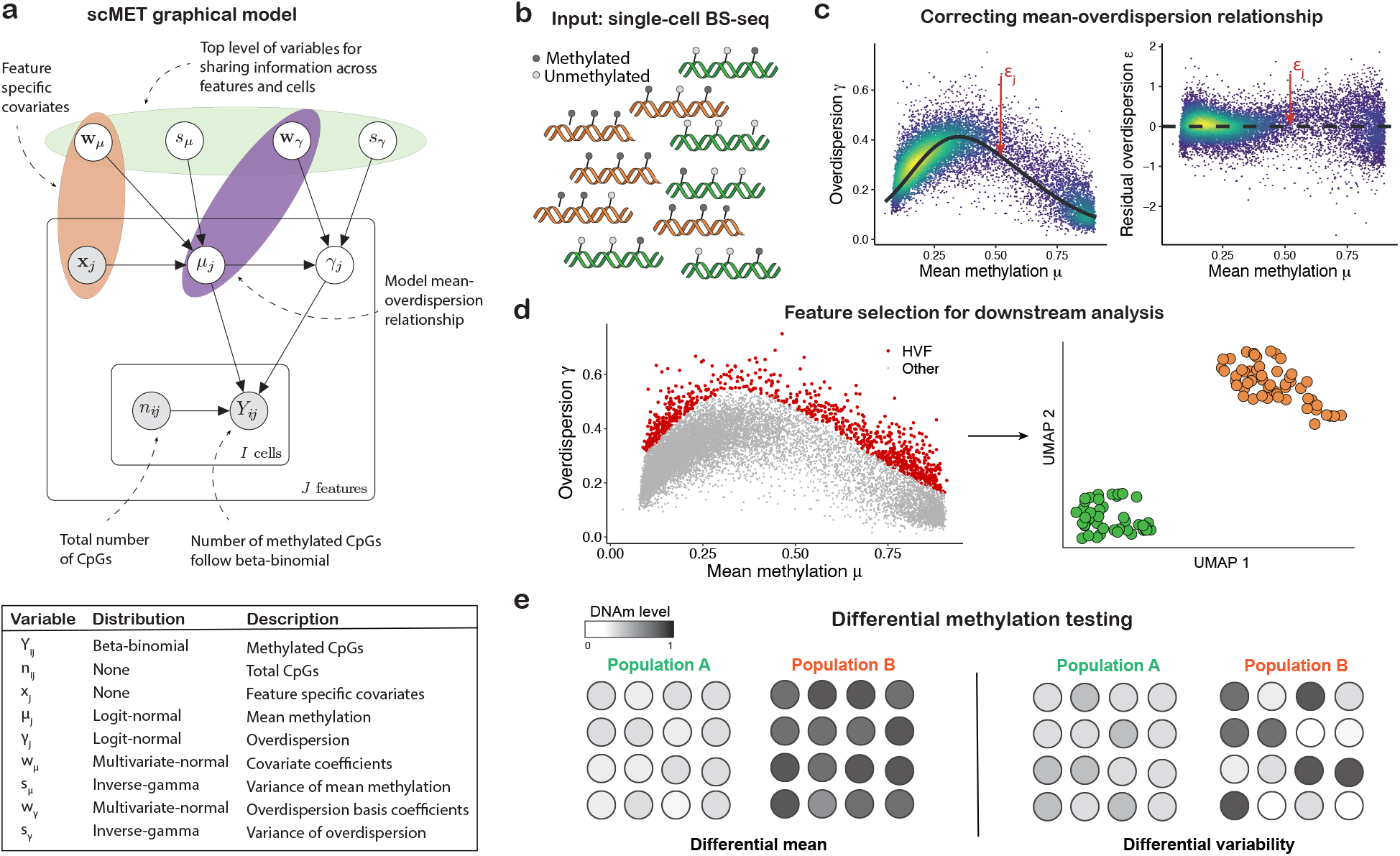
Graphical outline for scMET. (**a**) Overview of the scMET probabilistic graphical model. The random variables and data that form the model, along with the distributional assumptions, are shown. Input values are denoted by gray circles. Model parameters are denoted by white circles. (**b**) scMET uses single-cell DNAm data as input. The data could consist of measurements obtained from different groups of cells, such as experimental conditions or cell types (represented by green and orange colours in the diagram). For each region of interest (e.g. promoters), the input data is recorded in terms of the number of CpG sites for which a valid measurement was recorded and, among those, the number of methylated CpG sites. (**c**) By combining a hierarchical beta-binomial specification with a generalized linear model framework, scMET captures the mean-overdispersion relationship (left) that is typically observed in bisulphite sequencing readouts and derives residual overdispersion estimates that are not confounded by mean methylation (right). (**d**) scMET can be used to identify HVFs that drive epigenetic heterogeneity within a cell population. For example, these could be used as the input of dimensionality reduction techniques or clustering analyses. (**e**) scMET uses a probabilistic decision rule to perform differential methylation analysis: to identify features that show differences in mean methylation (left) and/or methylation variability (right) between pre-specified groups of cells.

The GLM framework is incorporated at two levels. Firstly, to introduce feature-specific covariates **x**_*j*_ (e.g. CpG density) that may explain differences in mean methylation *μ_j_* across features. Secondly, similar to Eling *et al.* (2018), we use a non-linear regression framework to capture the mean-overdispersion trend that is typically observed in high throughput sequencing data, such as scBS-seq (Fig. 1c). Critically, this trend is used to derive *residual overdispersion* parameters *ϵ*_*j*_ — a measure of cell-to-cell variability that is not confounded by mean methylation. Feature-specific parameters are subsequently used for: (i) feature selection, to identify highly variable features (HVFs) that drive cell-to-cell epigenetic heterogeneity (Fig. 1d) and (ii) differential methylation testing, to highlight features that show differences in DNAm mean or variability between specified groups of cells (Fig. 1e).

By using a Bayesian formulation, scMET infers the posterior distribution for all model parameters (*Methods*). Moreover, a variational Bayes scheme (Blei *et al.*, 2017) permits scalable analysis to thousands of cells and features (Supplementary Fig. S1), while having comparable posterior inference performance when compared to a Markov Chain Monte Carlo implementation (Supplementary Fig. S2 and S3). As in Bochkina and Richardson (2007), the output generated by scMET is used to implement a probabilistic decision rule to enable HVF selection and differential methylation testing. The decision rule is calibrated to control the expected false discovery rate (EFDR, Newton *et al.*, 2004). A more detailed description of the model specification and its implementation is provided in the *Methods* section. scMET is implemented as an R package and is available at https://github.com/andreaskapou/scMET.

### Benchmarking scMET on synthetic data

First, we benchmark the performance of scMET using synthetic data. To mimic the properties observed in real scBS-seq data, we simulated features with rich and poor CpG density (see *Methods* for details about the simulation settings). We compared mean and overdispersion estimates obtained by scMET with respect to BB maximum likelihood estimates (BB MLE), which were obtained separately for each feature. As expected, mean parameters *μ_j_* are easier to infer and estimates were comparable across both methods (Supplementary Fig. S4). However, scMET outperformed BB MLE when inferring overdispersion parameters *γ_j_*, particularly for small numbers of cells (Supplementary Fig. S5).

To assess whether the shrinkage introduced by scMET improves overdispersion estimates in real data, we performed down-sampling experiments based on a subset of the dataset introduced by *Luo et al.* (2017). For this analysis, we focused on 424 inhibitory neurons (a more detailed description is provided in *Methods*). We compared estimates obtained using the full and down-sampled datasets (Supplementary Fig. S6). We observed scMET posterior estimates to be more stable than BB MLE as the sample size decreased, suggesting that scMET leads to more robust inference. This is particularly important for rare cell populations or during early development, where large numbers of cells are difficult to obtain. In combination with the simulation study described above, this showcases the benefits of using a Bayesian hierarchical framework to share information across cells and genomic features.

Finally, we evaluated the performance of scMET as a tool to identify differentially methylated (DM) and differentially variable (DV) features. For this purpose, we generated synthetic data representing two cell types, for varying number of cells and different effect sizes in terms of log-odds ratio (Supplementary Note S2.3). For the DM analysis, we compared scMET against the Fisher’s exact test, which has been previously used for BS-seq data (e.g. Challen *et al.*, 2012). We observed better performance for scMET in terms of the F1-measure, especially for CpG rich features (Supplementary Fig. S7). scMET led to better type I error control, but to more conservative results (Supplementary Fig. S8). In terms of DV testing, the simulation study showed that for small effect sizes we would need more than 200 cells to achieve 50% to 80% power, whereas for features with larger effect sizes we would need around 50 cells per group to achieve 80% power (Supplementary Fig. S9).

### scMET improves feature selection for unsupervised analysis of single-cell methylomes from the mouse frontal cortex

To demonstrate the performance of scMET in real data, we considered a dataset where DNA methylation was profiled in 3,069 cells isolated from the frontal cortex of young adult mice (Luo *et al.*, 2017). To date, this is one of the largest and most heterogeneous publicly available scBS-seq datasets. The main source of heterogeneity in this dataset is due to two broad classes of neurons: excitatory (I = 2,645) and inhibitory (I = 424). Within each class, a hierarchy of sub-populations can be identified according to the cortical depth (Fig. 2a), where excitatory neurons progress from deep layers (mDL-1, mDL-2, mDL-3, mL6-1, mL6-2) to middle (mL5-1, mL5-2, mL4) and superficial layers (mL2/3). These groups were validated in the original study, and thus can be used as a benchmark for clustering analyses.

**Figure 2:**
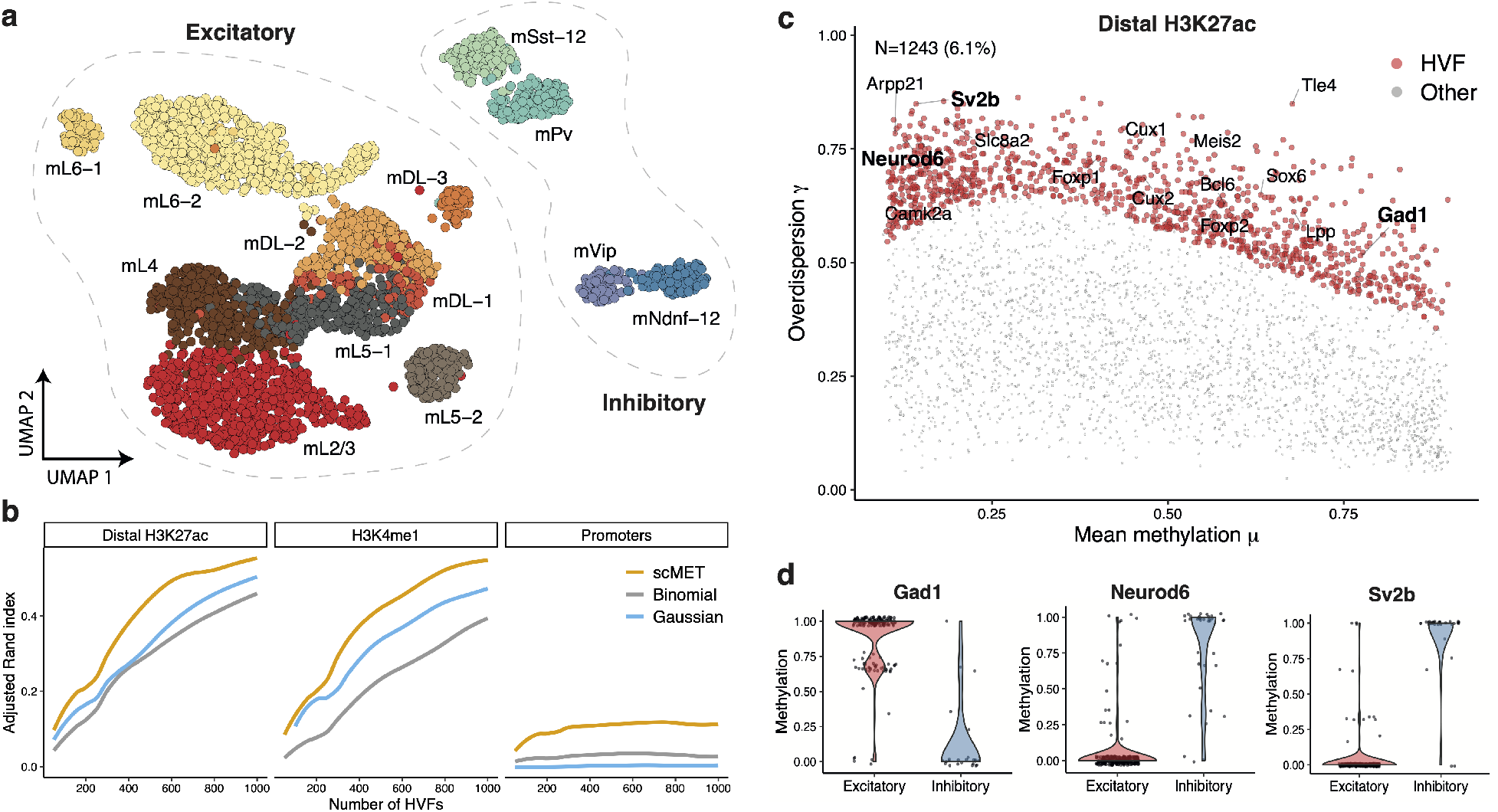
Feature selection using scMET characterises known heterogeneity on the mouse frontal cortex. (**a**) UMAP (Becht *et al.*, 2019) representation of neuron sub-populations present in the mouse frontal cortex dataset by combining the top 4,000 HVFs identified by scMET across distal H3K27ac and H3K4me1 genomic contexts. Cells are coloured according to the original cell sub-population assignments obtained by Luo *et al*. (**b**) Clustering performance in terms of adjusted Rand index (ARI), for varying number of HVFs. HVF selection was based on scMET’s residual overdispersion parameters *ϵ*_*j*_ (yellow), binomial variance (grey) and Gaussian variance (blue). A finite grid of HVFs was used for ARI evaluation and non-parametric regression was used to obtain a smoothed interpolation across all values (*Methods*). (**c**) Identifying HVFs for the distal H3K27ac genomic context. Red points correspond to features being called as HVF (EFDR = 10% and percentile threshold *δ_E_* = 90%). To ease interpretation each element is linked to its nearest gene. Labels highlight known neuron marker genes that were used in the Luo *et al.* (2017) study to define the different cell populations. (**d**) Example distal H3K27ac HVFs whose methylation patterns distinguish the two broad neuronal populations. Panel titles correspond to the nearest genes.

We applied scMET to genomic features from three different putative regulatory elements: gene promoters within 2kb around transcription start site (J = 12,774), distal H3K27ac peaks (J = 17,284) and H3K4me1 peaks (J = 30,374). As expected, scMET captured the mean-overdispersion relationship within each genomic context, and estimates for residual overdispersion parameters *ϵ*_*j*_ were not confounded by mean DNAm (Supplementary Fig. S10).

Here, we illustrate scMET as a feature selection tool, using residual overdispersion estimates to identify HVFs that can be used as input for unsupervised analyses, such as clustering. For each genomic context, we selected HVFs (Supplementary Fig. S11a) and performed a clustering analysis with varying numbers of HVFs (ranked by decreasing values of their associated tail posterior probabilities) as input. More concretely, we performed dimensionality reduction followed by *k*-means clustering (*Methods*) and used the adjusted Rand index (ARI, Hubert and Arabie, 1985) to quantify agreement with respect to the sub-populations validated by Luo *et al*. As a comparison, we also evaluated two alternative HVF selection strategies based on Gaussian and binomial models (*Methods*). As expected, the clustering performance improved steadily with increasing number of HVFs for all three methods. However, scMET consistently led to better clustering performance (Fig. 2b and Supplementary Fig. S11b, as well as Supplementary Fig. S12 and S13 for visual inspection in a low dimensional space). We could already separate inhibitory from excitatory neurons using only the top 100 HVFs obtained by scMET, and generally resulted in more distinct cell sub-populations. In all cases, promoters were less able to disentangle the neuronal sub-populations. This is consistent with the lower overdispersion levels observed in this genomic context (Supplementary Fig. S11c).

To facilitate interpretation for the HVFs highlighted by scMET, we linked genomic features to genes by overlapping the genomic coordinates allowing for a maximum distance of 20kb from the transcription start site in the case of distal elements. We explored whether features identified as HVF (red points in Fig. 2c and Supplementary Fig. S14a) were enriched for neuronal markers identified in the *Luo et al.* (2017) study (Supplementary Table S1). This enrichment was observed for distal H3K27ac and H3K4me1 marks, but not for promoters (Supplementary Fig. S14b). As representative examples, we display three distal H3K27ac elements among the HVFs that are located proximal to known gene markers of each neuron class: *Gad1* for inhibitory, *Neurod6* and *Sv2b* for excitatory (Fig. 2d).

### scMET enables differential methylation testing between cellular sub-populations

To showcase scMET as a differential methylation tool, we applied it on the same mouse frontal cortex dataset (Luo *et al.*, 2017), after separating the cells in excitatory and inhibitory groups. Initially, we applied scMET to characterise differential methylation (DM), i.e. changes in mean methylation. Across all genomic contexts, we observed a substantially larger fraction of features being hyper-methylated in inhibitory compared to excitatory neurons (Fig. 3a and Supplementary Fig. S15a). Within distal H3K27ac peaks, for instance, scMET identified 5,242 features to have higher methylation levels in inhibitory neurons, compared to only 935 features showing higher methylation in the excitatory group (Fig. 3a-b). After mapping features to their nearest gene, we observed that DM hits were enriched for known marker genes that differentiate inhibitory and excitatory neurons (Supplementary Fig. S15b).

**Figure 3:**
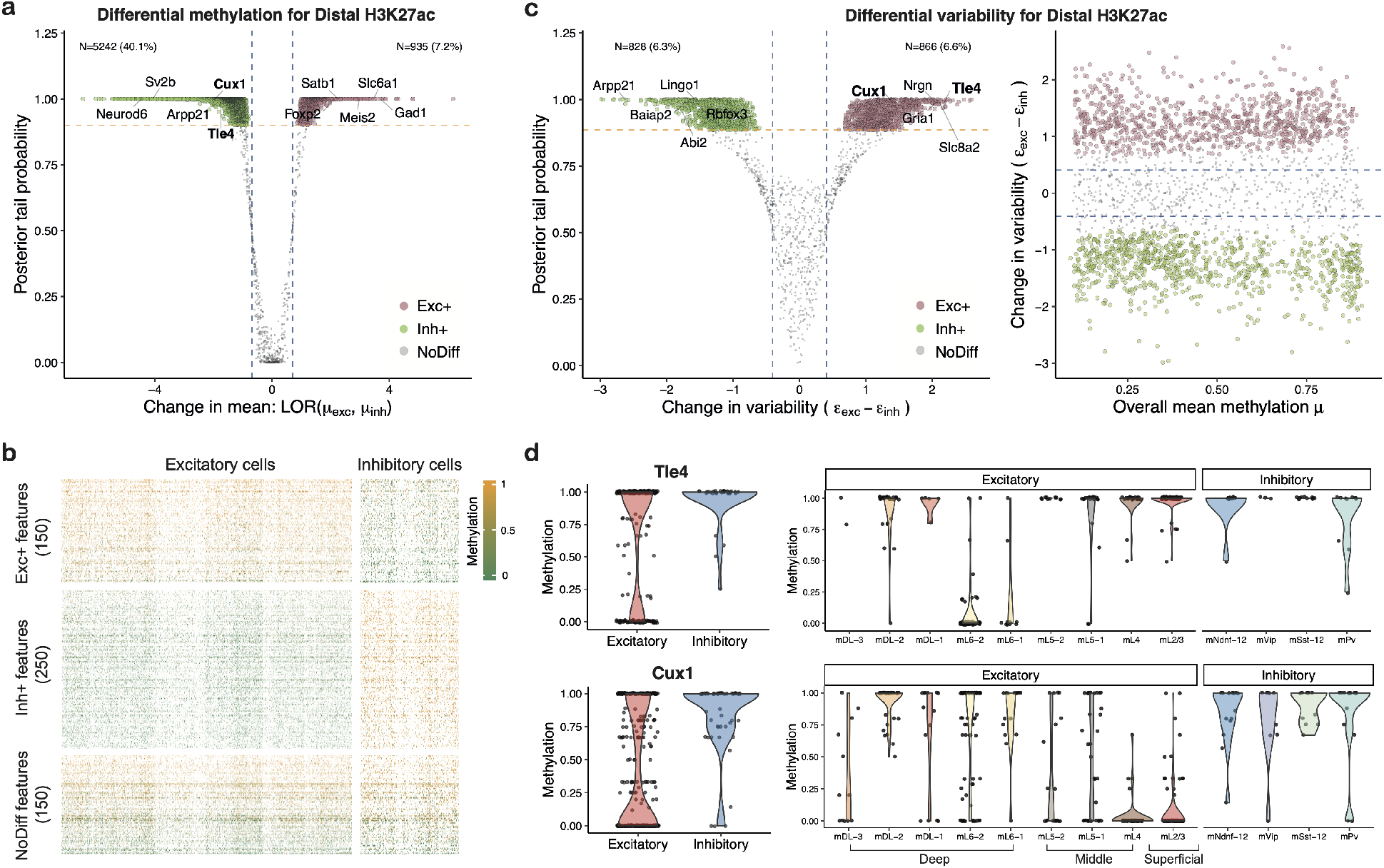
Summary of changes in methylation patterns (mean and variability) for inhibitory and excitatory neurons. (**a**) Identifying differentially methylated (DM) features for the distal H3K27ac genomic context. Green and pink points correspond to features showing higher mean methylation in inhibitory (Inh+) and excitatory (Exc+) neurons, respectively. Labels highlight neuron marker genes that were used in Luo *et al.* (2017) to define the different cell populations. Blue vertical dashed lines correspond to log-odds ratio threshold *ψ*_*M*_ = ± log(2). Yellow horizontal dashed line is located at posterior evidence probability cut-offs defined by EFDR = 5%. (**b**) Representative heatmap of methylation rates (*Y*_*ij*_/*n*_*ij*_) across cells (columns) and features (rows) for the distal H3K27ac genomic context. Cells are grouped in excitatory and inhibitory classes. A set of randomly selected features is displayed. These are grouped according to the DM analysis output as: Exc+, Inh+ and no mean methylation difference. The colour code represents features with low (0, green) and high (1, yellow) mean methylation level. Features with no CpG coverage are denoted with white colour. (**c**) Identifying differentially variable (DV) features for the distal H3K27ac genomic context. Green and pink points correspond to features showing higher methylation variability in inhibitory and excitatory neurons, respectively. Blue dashed lines correspond to log-odds ratio threshold *ψ_E_* = ± log(1.5). Yellow dashed line is located at posterior evidence probability cut-offs defined by EFDR = 5%. For each feature, posterior estimates for the change in residual overdispersion parameter *ϵ*_*j*_ between excitatory and inhibitory neurons is plotted against the posterior tail probability of calling a feature as DV (left). For each feature, posterior estimates for mean methylation parameter *μ*_*j*_ is plotted against posterior estimates for the change in residual overdispersion parameter *ϵ*_*j*_ between excitatory and inhibitory neurons (right). (**d**) Example features that are called as being more variable in excitatory neurons. Left subplots show broad differences in methylation patterns. Right subplots show methylation patterns separately within each broad neuronal class. Each data point represents the methylation rate for a cell.

Besides DM testing, the primary focus of the scMET differential test is to identify changes in cell-to-cell methylation variability. In principle, differential variability (DV) testing could be based on feature-specific overdispersion parameters *γ_j_*, but these results would be confounded by the mean-overdispersion trend (Fig. 1c). Hence, meaningful DV analysis based on *γ_j_* would need to be restricted to non-DM features. Instead, we propose to perform DV analysis based on residual overdispersion parameters *ϵ*_*j*_. For the mouse frontal cortex dataset, we identified a large number of DV features across genomic contexts, except from promoter regions which showed tighter methylation patterns across inhibitory and excitatory neurons (Fig. 3c and Supplementary Fig. S16a). Critically, the procedure for calling DV features was not confounded by mean methylation levels (Fig. 3c and Supplementary Fig. S16b).

As representative examples, we show two distal H3K27ac peaks that are located proximal to neuronal markers and exhibit higher variability in excitatory neurons (Fig. 3d). Both features show substantial variation across the different sub-types of excitatory neurons: the first is mostly unmethylated in the mL6 cortical layer, the second is mostly unmethylated in the superficial cortical layer. These patterns are consistent with previously reported spatial expression for *Tle4* (mostly expressed in the deep cortical layer, see Sorensen *et al.*, 2015; Zahr *et al.*, 2018) and *Cux1* (which shows expression specificity for the superficial layer, see Georgala *et al.*, 2011; Zahr *et al.*, 2018). It should be noted that these features would be excluded from DV analysis based on *γ_j_*, since they are also DM between the two broad classes (Fig. 3a). In summary, these findings demonstrate the ability of scMET to identify potential markers that drive between and within cell population heterogeneity.

### Exploring the relationship between transcriptional and DNAm variability using single-cell multi-omics data

As a second use case, we considered a single-cell multi-omics dataset where scNMT-seq (Clark *et al.*, 2018) was employed to profile RNA expression, DNAm and nucleosome occupancy at single-cell resolution, spanning multiple time points from the exit from pluripotency to primary germ layer specification (Argelaguet *et al.*, 2019). The multi-modal nature of this dataset provides a unique opportunity to link cell-to-cell variation between DNAm and transcription across individual cells. Here we used scMET to quantify DNAm variability at promoter elements, which we subsequently contrasted to RNA expression heterogeneity for the corresponding genes. For this analysis we exclusively used promoter elements as, unlike distal regulatory elements, they can be unambiguously matched to their respective genes.

For each gene, we quantified transcriptional heterogeneity using the residual overdispersion estimates generated by BASiCS (Supplementary Fig. S17a, Eling *et al.*, 2018). Promoter DNAm variability was calculated using the residual overdispersion estimates inferred using scMET (Supplementary Fig. S17b). More details about these analyses and the associated data pre-processing steps are described in the *Methods* section.

When comparing residual overdispersion estimates for RNA expression and promoter DNAm, there was no clear genome-wide association (Fig. 4a). However, when restricting to genes that display high levels of transcriptional variability, two main groups can be identified. The first category corresponds to genes with low levels of promoter DNAm residual overdispersion, and it includes differentiation and germ layer markers such as *Mesp1*, *Lefty2*, *Id3* (mesoderm) and *Cldn6*, *Cer1* and *Krt8* (endoderm). The second category is characterised by genes with high promoter DNAm residual overdispersion and includes known pluripotency markers such as *Dppa5a*, *Zfp42*, *Spp1* and *Peg3*. Representative examples for these genes are displayed in Fig. 4b.

**Figure 4:**
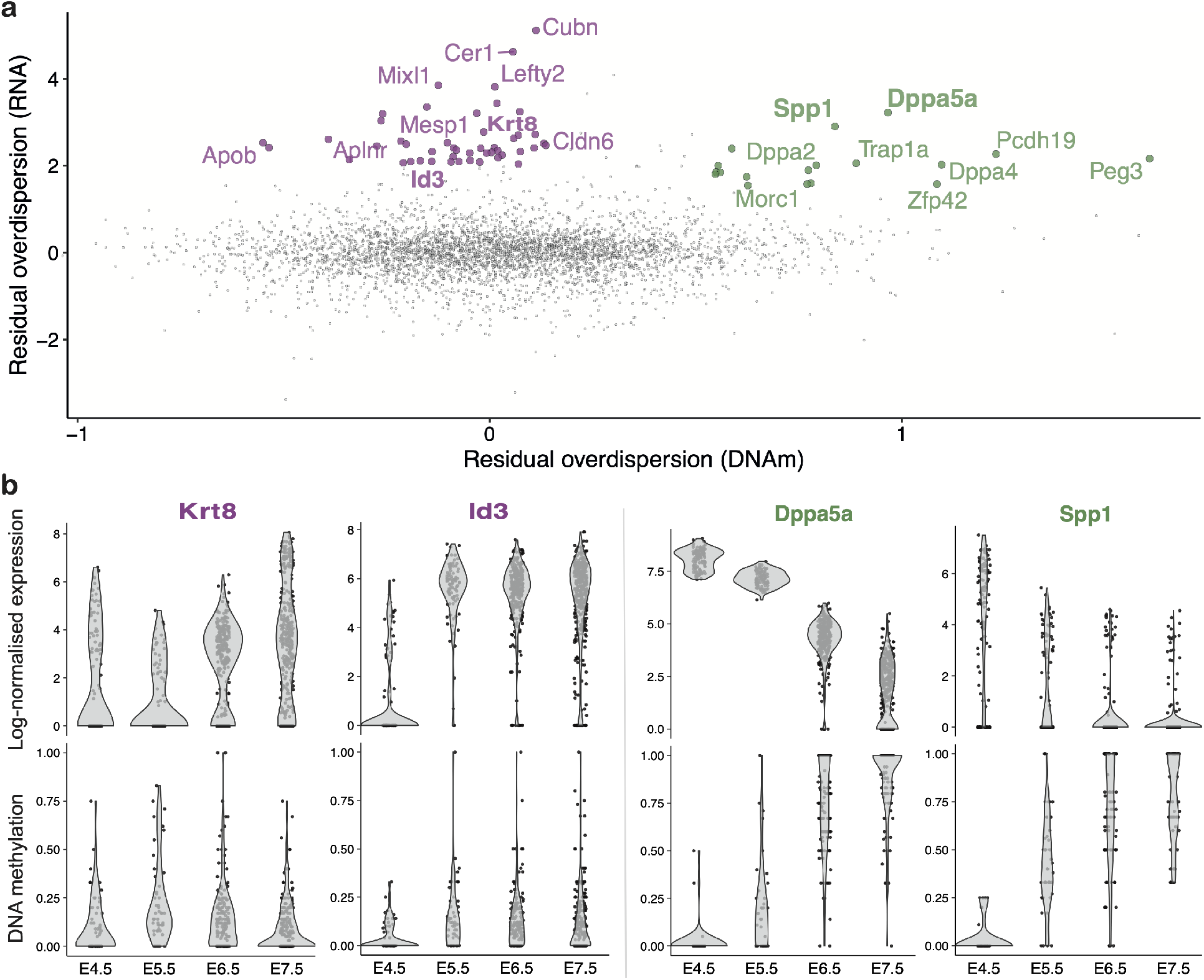
scMET applied to the multi-omics scNMT-seq gastrulation dataset reveals a complex linkage between promoter DNAm and RNA expression during embryonic development. (**a**) Scatter plot displays posterior median estimates for residual DNAm overdispersion parameters *ϵ*_*j*_ in gene promoters (x-axis) versus RNA residual overdispersion of the corresponding genes (y-axis). Among the genes with high levels of RNA heterogeneity, green and pink points correspond to promoters showing high and low levels of DNAm variability, respectively. (**b**) Representative examples of DNAm and RNA expression patterns across developmental stages for genes with high transcriptional heterogeneity and low (left, pink) or high (right, green) DNAm heterogeneity. Y-axis shows BASiCS log-normalised gene expression (in a log(*x* + 1) scale) (top) and promoter DNAm rate (bottom). Cells are stratified by embryonic stage (x-axis).

These results suggest the presence of two modes of regulation. On one hand, down-regulation of pluripotency genes is associated with high promoter DNAm heterogeneity, linked to a pronounced increase in promoter DNA methylation throughout the embryonic stages. On the other hand, up-regulation of differentiation genes is not linked to high levels of promoter DNAm variability. This suggests that other genomic contexts or molecular layers might be responsible for their activation (Argelaguet *et al.*, 2019). Finally, we also find genes with low RNA expression variability that display high levels of promoter DNAm heterogeneity (Supplementary Fig. S17c), suggesting that the coupling between promoter DNAm and transcriptional activity is more complex than previously acknowledged during embryonic development stages (Anastasiadi *et al.*, 2018).

## Discussion

Single-cell DNAm assays can currently profile hundreds to thousands of DNA methylomes, with increasingly complex experimental designs. The high resolution of these measurements enables us to measure cell-to-cell epigenetic variability, as well as uncover the regulatory features that modulate it (Eling *et al.*, 2019). However, the noise and biases intrinsic to such technologies create a need for computational frameworks that can systematically interrogate the data generated, dissecting genuine variability and quantifying uncertainties.

In this study we introduced scMET, a statistical framework for modelling DNA methylation heterogeneity from scBS-seq data. Using a hierarchical Bayesian framework to borrow information across cells and features, scMET robustly quantifies genuine cell-to-cell variability. Our results demonstrated the ability of scMET in highlighting genomic features that drive cell-to-cell heterogeneity across neuronal sub-populations in a large dataset of single-cell methylomes from the mouse frontal cortex. Furthermore, scMET can be used as a quantitative tool to interrogate changes in DNAm patterns between pre-specified cell populations. Unlike common approaches that only detect changes in mean methylation levels (Feng *et al.*, 2014; Hansen *et al.*, 2012), scMET can also identify features with differences in DNAm variability between populations. Importantly, the differential variability estimates are quantified through residual overdispersion parameters, thus accounting for the known confounding relationship between mean and overdispersion in scBS-seq datasets.

scMET uses a GLM framework to explicitly model known biases in the data in the form of additional covariates, such as CpG content. The flexibility of the GLM approach enables it to easily incorporate additional features, such as DNA motifs, which could be important to elucidate the role of sequence or chromatin state in modulating DNA methylation. Additionally, the framework could readily be extended to model joint variability in multiple molecular layers (such as transcriptome and methylome), opening a path to new methodologies in integrative, single-cell multi-omics analyses. Given the increasing prominence of such studies, we expect scMET to become an important tool in the extraction of biological signals from DNAm datasets of increasing complexity.

## Methods

Standard statistical models for count data, such as the Poisson and binomial distributions, do not always capture the properties of data generated by high-throughout sequencing assays (e.g. RNA sequencing, bisulphite sequencing). In such cases, the data typically exhibit higher variance than what is predicted by these models — this is often referred to as *overdispersion* (Cox, 1983; Hinde *et al.*, 1998). This overdispersion may relate to *technical variation* (e.g. due to differences in sequencing depth) or to *biological variation* between the units of interest (e.g. cells or subjects) that is linked to genetic, environmental or other factors. Disentangling these sources of variation is a major challenge in computational biology.

### The scMET model

Let *Y_ij_* represent the number of methylated CpGs out of the *n_ij_* CpGs for which DNAm was measured for genomic feature *j* ϵ {1,…, *J*} in cell *i* ϵ {1*,…, I*}. These genomic features could be defined by pre-annotated regions (e.g. enhancers) or other regions of interest. To capture data overdispersion, scMET assumes a beta-binomial (BB) hierarchical formulation:

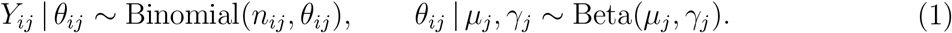

In Eq. (1), the beta distribution is parameterised such that 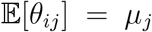 and 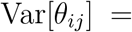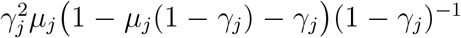, with *μ_j_* ∈ (0, 1) and *γ_j_* ∈ (0, 1). If *γ_j_* = 0, the model in Eq. (1) reduces to a binomial model with parameters *n_ij_* and *μ_j_*. After integrating out the random effects *θ_ij_*, it can be seen that *μ_j_* corresponds to the mean methylation across all cells for feature *j* and that *γ_j_* controls the *overdispersion* that is not captured by binomial sampling. In fact, the BB variance can be written as:

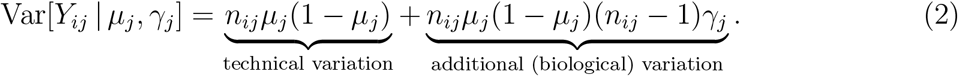

Parameters *μ_j_* and *γ_j_* can be inferred via maximum likelihood estimation. However, due to the high sparsity and noise present in single-cell DNAm data, these estimates can be unstable, especially for overdispersion parameters *γ_j_* (Supplementary Fig. S5 and S6). To overcome this, we use a Bayesian framework with a hierarchical prior specification for *μ_j_* and *γ_j_*, sharing information across sets of similar types of genomic features (e.g. enhancers). Our approach is flexible and can incorporate feature-specific covariates **x**_*j*_ that explain differences in mean methylation across features. For instance, features with high CpG density tend to have lower methylation levels. These covariates are introduced within a generalized linear model (GLM) framework through the prior on the mean methylation parameters *μ_j_*:

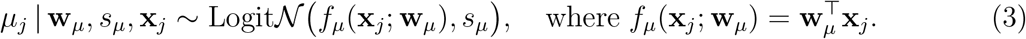

In Eq. (3), 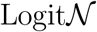 denotes a logit-normal distribution, **w**_*μ*_ is a vector of regression coefficients and *s_μ_* is the standard deviation for logit(*μ_j_*). Throughout our analyses we assume **x**_*j*_ = (1*, C_j_*), where *C_j_* denotes the CpG density for feature *j*. However, scMET is flexible and users can introduce other feature-specific covariates.

Our prior specification is also designed to capture the mean-overdispersion relationship that is typically observed in the data generated by high-throughput sequencing assays, such as scBS-seq (Fig. 1c). Here, we follow the approach in *Eling et al.* (2018), introducing a non-linear regression model through an informative prior for *γ_j_*:

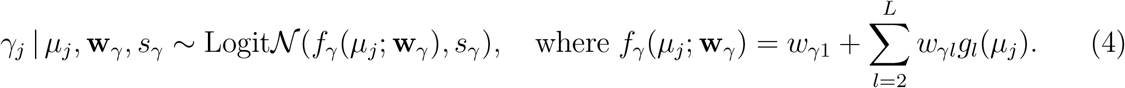

Here, *f_γ_*(*μ_j_*; **w**) can be interpreted as the overdispersion (logit scale) that is predicted by mean methylation levels *μ_j_* (fitted black line in Fig. 1c), *g_l_*(*μ_j_*) represent radial basis function kernels (defined as in Kapourani and Sanguinetti, 2016), and *w_γ_*_1_*,…, w_γ,L_* are regression coefficients. Unless otherwise stated, we use *L* = 4 throughout our analyses. The remaining elements of the prior are described in Supplementary Note 2.1.

The prior distribution in Eq. (4) can be rewritten as a non-linear regression model

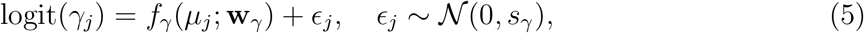

where *ϵ*_*j*_ corresponds to a feature-specific *residual overdispersion* parameter that captures deviations from the overall trend. Hence, a feature that exhibits positive *ϵ*_*j*_ values has more variation than expected for features with similar mean methylation. Accordingly, negative *ϵ*_*j*_ values indicate less variation than expected for features with similar mean methylation.

### Implementation

The posterior distribution for the model parameters in scMET is not amenable to analytic solutions. Hence, we resort to variational Bayes (VB, Blei *et al.*, 2017) and Markov Chain Monte Carlo (MCMC, Liang *et al.*, 2010) implementations using the Stan probabilistic programming language (Carpenter *et al.*, 2017). scMET is publicly available as an R package at https://github.com/andreaskapou/scMET and will be shortly submitted to Bioconductor.

### Identifying highly variable features

Residual overdispersion parameters *ϵ*_*j*_ can be used to label highly variable features (HVFs) within a population of cells. Our decision rule is based on tail posterior probabilities (Bochkina and Richardson, 2007) associated to whether *ϵ*_*j*_ exceed a pre-specified threshold *ϵ*_0_:

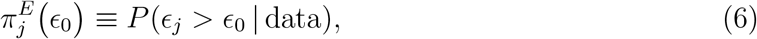

As a default choice, we define *ϵ*_0_ based on the distribution of posterior estimates for residual overdispersion parameters *ϵ*_*j*_ across all features. In particular, we define *ϵ*_0_ to match the *δ_E_*-th percentile of the distribution. Unless otherwise stated, we set as default *δ_E_* = 0.9.

The probabilities in Eq. (6) can be estimated by counting the proportion of posterior draws (obtained by VB or MCMC) for which the chosen criteria are met (Lewin *et al.*, 2006). scMET labels as HVFs those for which their associated tail posterior probabilities are above a given posterior evidence threshold *α_H_* (0.6 < *α*_*H*_ < 1), where *α_H_* is calibrated via the expected false discovery rate (EFDR; Newton *et al.*, 2004), see also Supplementary Note S2.2.

### Differential testing

scMET provides a similar probabilistic rule to label differentially methylated (DM) and differentially variable (DV) features across experimental conditions or cell types (Fig 1e). Here, we define DM features as those for which mean methylation varies across the groups of cells under study. More concretely, let 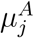 and 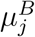 be the mean methylation parameters associated with feature *j* in groups A and B. We quantify differences in mean methylation as the log-odds ratio (LOR):

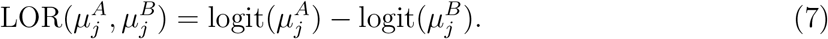

Similar to the HVF analysis, our decision rule for DM testing is defined as:

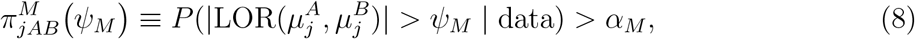

where *α_M_* (0.6 < *α*_*M*_ < 1) is a posterior evidence threshold chosen to match a desired EFDR level and *ψ_M_* is a LOR threshold which can be interpreted as a minimum effect size to be detected by the test. As default, we use *ψ_M_* = log(2), i.e. a two fold change in odds ratio.

Beyond highlighting DM features, scMET embraces the cellular resolution of scBS-seq data to perform differential variability (DV) analyses, identifying changes in cell-to-cell DNAm variability across groups. Although overdispersion parameters *γ_j_* could be used as the input for the DV test, the results would be confounded by the mean-overdispersion relationship that is typically observed within each genomic context (Fig 1c). Instead, we propose to perform DV analysis based on *ϵ*_*j*_ — a measure of cell-to-cell DNAm variability that is not confounded by differences in mean methylation. Let 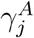 and 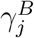 denote the overdispersion parameters linked to feature *j* in groups A and B. To label DV features based on residual overdispersion, we make use of Eq. (5), and decompose the LOR between 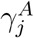 and 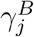 parameters as:

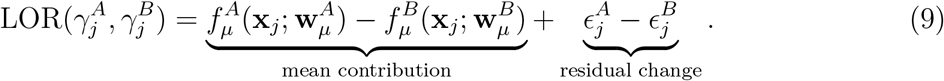

In Eq. (9), the first term captures the changes in overdispersion that are explained by mean methylation and the second term captures residual overdispersion changes after accounting for the mean methylation. This residual change is used to identify features with statistically significant differences in residual overdispersion. For a given posterior evidence threshold *α_E_* (0.6 < *α*_*E*_ < 1) and tolerance threshold *ψ*_*E*_, the following rule is used to identify DV features:

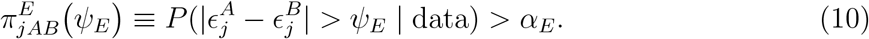

As default we set *ψ_E_* = log(1.5), i.e. 50% change in overdispersion LOR between the groups. As above, the posterior evidence threshold *α_E_* is calibrated via the EFDR, see Supplementary Note 2.2.

### Competing methods

The BB MLE method corresponds to estimating the parameters of the beta-binomial model in Eq. (1) independently per feature using maximum likelihood. The VGAM package was used for parameter estimation (Yee, 2015).

For HVF selection, two additional strategies were considered. The binomial model, where features are ranked according to binomial variance given by 1/*I Σ*_*i*_ *θ*_*ij*_(1 − *θ_ij_*), where *θ_ij_* = *Y_ij_/n_ij_* is the methylation rate for feature *j* in cell *i*. The Gaussian model on methylation rates 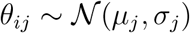, where features are ranked according to *σ_j_*.

For differential mean methylation testing on the synthetic datasets, we compared scMET with the Fisher’s exact test. Features with log-odds ratio > log(1.5) between the two groups and FDR < 10% (based on Benjamini-Hochberg procedure) were called as differentially methylated. Whilst methods have been proposed for DV testing using bulk methylation data (e.g. Phipson and Oshlack, 2014), to the best of our knowledge, scMET is the first DV approach tailored to single cell methylation data.

### Simulation study

We simulated J = 300 features for varying number of cells ranging from I = 20 up to I = 1000. To mimic the properties observed in real scBS-seq data, we assume that for each feature we have coverage for a subset of cells given by *I_j_* ∼ Binomial(*I, p_j_*), where *p_j_* ∼ Uniform(0.4, 0.8) to generate diverse *I_j_* across features. We also simulate two alternative regions that have rich (N = 50) and poor (N = 15) CpG density. That is, the number of CpGs (*n_ij_*) are simulated from Binomial(*N, q_j_*), where *q_j_* ∼ Uniform(0.4, 0.8) to generate a broad range of CpG coverage across features. Next, for each feature we generate mean methylation parameters 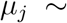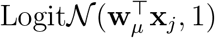 where **w**_*μ*_ = (−0.5, −1.5) and **x**_*j*_ = (1*, C_j_*) are feature-specific covariates, where *C_j_* denotes the CpG density. The negative weight on **w**_*μ*_ is used to simulate the known negative association between mean methylation and CpG density. Next, we simulated feature-specific overdispersion parameters 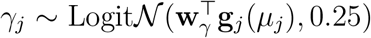 to mimic the mean-overdispersion relationship. We set **w**_*γ*_ = (−1.2, −.3, 1.1, −.9) and **g**_*j*_(*μ_j_*) is a vector of basis function values with methylation level *μ_j_*. Finally, we simulated the number of methylated CpGs from BB distribution, using the `VGAM` package, that is, *Y_ij_* ∼ BB(*n_ij_, μ_j_, γ_j_*).

For differential testing analysis, we used the above approach to generate cells from the first group (group A). For DM analysis, 15% of features were randomly selected and their corresponding *μ_j_* were randomly increased or decreased by three different LOR thresholds: 2, 3, and 5, to generate cells from the second group (group B). Similarly, for DV analysis 15% of features were randomly selected from the first group and their corresponding *γ_j_* were randomly increased or decreased by three different LOR thresholds: 2, 3, and 5, to generate cells from second group.

### Mouse frontal cortex dataset

#### Data processing

The dataset is available from the Gene Expression Omnibus repository under accession number GSE97179. Details on quality control and data pre-processing can be found in *Luo et al.* (2017). Supplementary Table S2 contains metadata for the 3,069 cells, such as cell type annotations and cortical layer information. We aggregated closely related cellular sub-populations with less than 25 cells, following the hierarchy established in Luo *et al.* (2017). DNA methylation was quantified using mCG dinucleotides over three genomic contexts: (1) gene promoters (±2kb windows around the transcription start sites of genes extracted from ENSEMBL version 87 Yates *et al.*, 2016), (2) Distal H3K27ac ChIP-seq peaks and (3) H3K4me1 ChIP-seq peaks. The latter was based on two ChIP-seq datasets that were profiled in adult (8 weeks) mouse cortex as part of the ENCODE project (see Supplementary Table S3).

For each genomic feature *j* ∈ {1,…, *J*} and cell *i* ∈ {1,…, *I*}, the following censoring procedure was applied: we recorded *Y_ij_* as a missing value if methylation coverage was available in less than 3 CpGs (i.e. *n*_*ij*_ < 3). The purpose of this censoring step was to exclude observations with very low coverage for which DNAm quantification is less reliable. Subsequently, we removed features that did not have CpG coverage in at least 15 cells. In addition, we excluded features that had mean methylation across cells lower than 0.1 or higher than 0.9; the rationale being that fully (un)methylated features do not drive methylation heterogeneity and will not provide information for identifying cell sub-populations.

#### Down-sampling experiment

Using the characterised sub-populations from Luo *et al.* (2017) we performed down-sampling experiments on 424 inhibitory neurons. Both scMET and BB MLE methods were run once on the full dataset (424 cells) to generate pseudo-ground truth parameter estimates. Subsequently, 20, 50, 100 and 200 cells were randomly down-sampled from the full population prior to parameter estimation. This procedure was repeated 5 times for each sample size. The same censoring step as described above was applied. Moreover, due to smaller sample sizes, we filtered genomic features that did not have CpG coverage in at least 5 cells.

#### HVF analysis

HVF analysis was applied on 12,774 gene promoters, 17,284 distal H3K27ac peaks, and 30,374 H3K4me1 peaks. To model the mean-overdispersion relationship we used L = 4 radial basis function kernels and kept default hyper-parameter values (Supplementary Note S2.1). The total number of iterations was set to 50,000 and convergence was attained when the evidence lower bound difference between two consecutive iterations was less than 1e-04.

#### Differential analysis

For differential analysis between excitatory and inhibitory neurons, we only included features with CpG coverage in at least 15 cells, in both sub-populations. This resulted in 12,611 gene promoters, 13,075 distal H3K27ac peaks, and 20,212 H3K4me1 peaks. To model the mean-overdispersion relationship we used L = 4 radial basis function kernels and kept default hyper-parameter values (Supplementary Note S2.1). The total number of iterations was set to 50,000 and convergence was attained when the evidence lower bound difference between two consecutive iterations was less than 1e-04.

#### Dimensionality reduction

Dimensionality reduction was applied using a Bayesian Factor Analysis algorithm, as implemented in the MOFA2 package (Argelaguet *et al.*, 2020). The motivation for this method, as opposed to the conventional Principal Component Analysis, is to handle the large presence of missing values without need for imputation. A second (non-linear) dimensionality reduction step was applied using UMAP (as implemented in the umap package) to project the data into a two-dimensional space (Supplementary Fig. S12 and S13).

#### Clustering

A finite grid of HVFs (from 50 to 1,000 with step size of 50) was selected by each of the competing methods. Subsequently, clustering analysis was performed using the *k*-means algorithm on the latent space defined by the MOFA factors (fixed to 15). The number of clusters was set to the number of cell types as characterised by *Luo et al.* (2017) (see also *Data processing* section). We assessed clustering performance using the ARI and cluster purity. A non-parametric regression (implemented by the loess function) was used to obtain a smoothed interpolation across all HVF values.

### scNMT-seq gastrulation dataset

#### Data processing

The parsed scNMT-seq gastrulation dataset was downloaded from ftp://ftp.ebi.ac.uk/pub/databases/scnmt gastrulation. Raw files are available from the Gene Expression Omnibus repository under accession number GSE121708. Details on the quality control and data processing can be found in Argelaguet *et al.* (2019). We selected all cells from E4.5 to E7.5 days after excluding the extra-embryonic visceral endoderm cells, as they display distinct DNA methylation profiles. Supplementary Table S4 contains sample metadata for the 848 cells retained for analysis. DNA methylation was quantified over gene promoters (±2kb windows around the transcription start sites of genes extracted from ENSEMBL version 87 Yates *et al.*, 2016).

#### Calculation of DNA methylation and RNA expression heterogeneity

For the DNAm data, we applied the same censoring procedure and feature exclusion criteria as described in the pre-processing of the Luo *et al.* (2017) dataset. This resulted in 13,785 gene promoters for downstream analysis. Residual overdispersion estimates were calculated by scMET with default parameter values using the same number of iterations and convergence criteria described above.

For the RNA expression data, we removed lowly expressed genes (no counts in less than 10 cells and average count across expressed cells less than 5). This resulted in 14,076 genes for downstream analysis. Residual overdispersion estimates were obtained using BASiCS (Eling *et al.*, 2018). The algorithm was run using 20,000 iterations, applying a burn in of 10,000 and thinning of 10. An empirical Bayes approach was used to derive the prior hyperparameters associated to gene-specific mean expression parameters within BASiCS.

In the comparison displayed in Fig 4, we focused on the 10,192 genes contained in the intersection of the lists obtained above.

## Data availability

All datasets analysed in this article are publicly available in the cited references.

## Code availability

scMET is implemented in an open-source and publicly available R package at https://github.com/andreaskapou/scMET. The code used to process and analyse the data is available at https://github.com/andreaskapou/scMET-analysis. The following software versions were used throughout our analyses: scMET (0.99.1), BASiCS (2.0.1), coda (0.19.3), MOFA2 (0.99.7), rstan (2.19.3) and VGAM (1.1.3).

## Acknowledgements

We thank Oliver Stegle and John C. Marioni for constructive feedback during early stages of this project. C.A.K. is a cross-disciplinary post-doctoral fellow supported by funding from the University of Edinburgh and Medical Research Council (core grant to the MRC Institute of Genetics and Molecular Medicine). C.A.V. is a Chancellor’s Fellow funded by the University of Edinburgh. R.A. was funded by the European Molecular Biology Laboratory (EMBL) international PhD programme.

## Authors’ contributions

All authors conceived and planned the methods and experiments. C.A.K. developed the statistical model and software implementation. C.A.K. and R.A. analysed the data and wrote the manuscript, with input from all authors. G.S. and C.A.V. supervised the study. All authors contributed to the interpretation of results and approved the final manuscript.

## Ethics approval and consent to participate

Ethical approval was not needed for this study.

