## Supplementary Figures and Notes for "scMET: Bayesian modelling of DNA methylation heterogeneity at single-cell resolution"

### S1 Supplementary figures

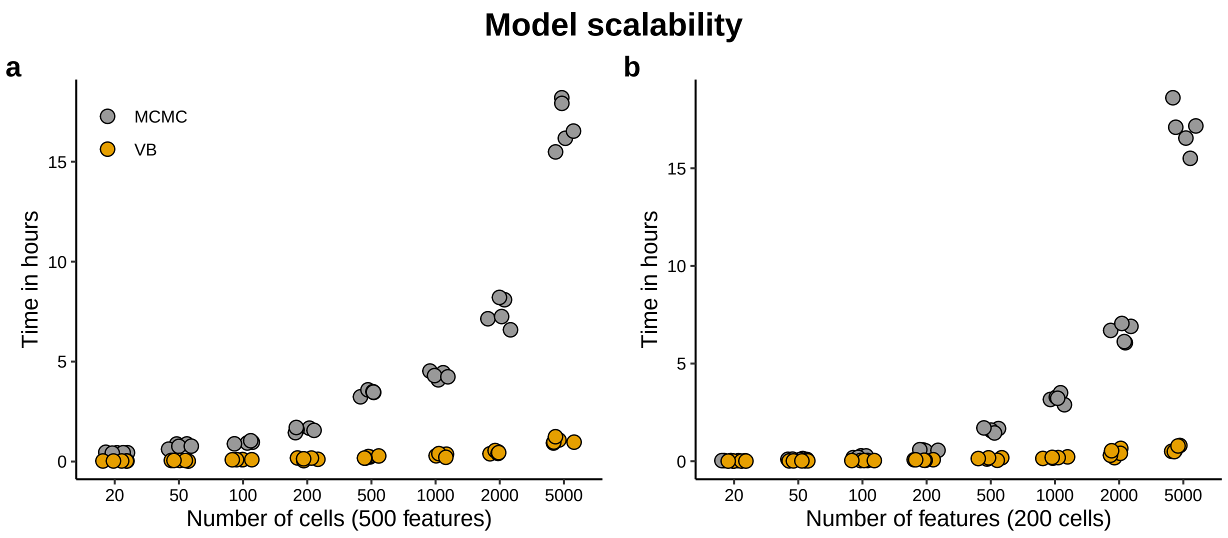


Figure S1: scMET scalability analysis using synthetic data. Running times for varying number of (**a**) cells and (**b**) features. Each dot represents a different run comparing variational Bayes (VB, yellow) and Markov Chain Monte Carlo (MCMC, grey) implementations of scMET in Stan (Carpenter *et al.*, 2017). Total number of iterations for VB was set to a maximum of 20,000, whereas for MCMC to 3,000.

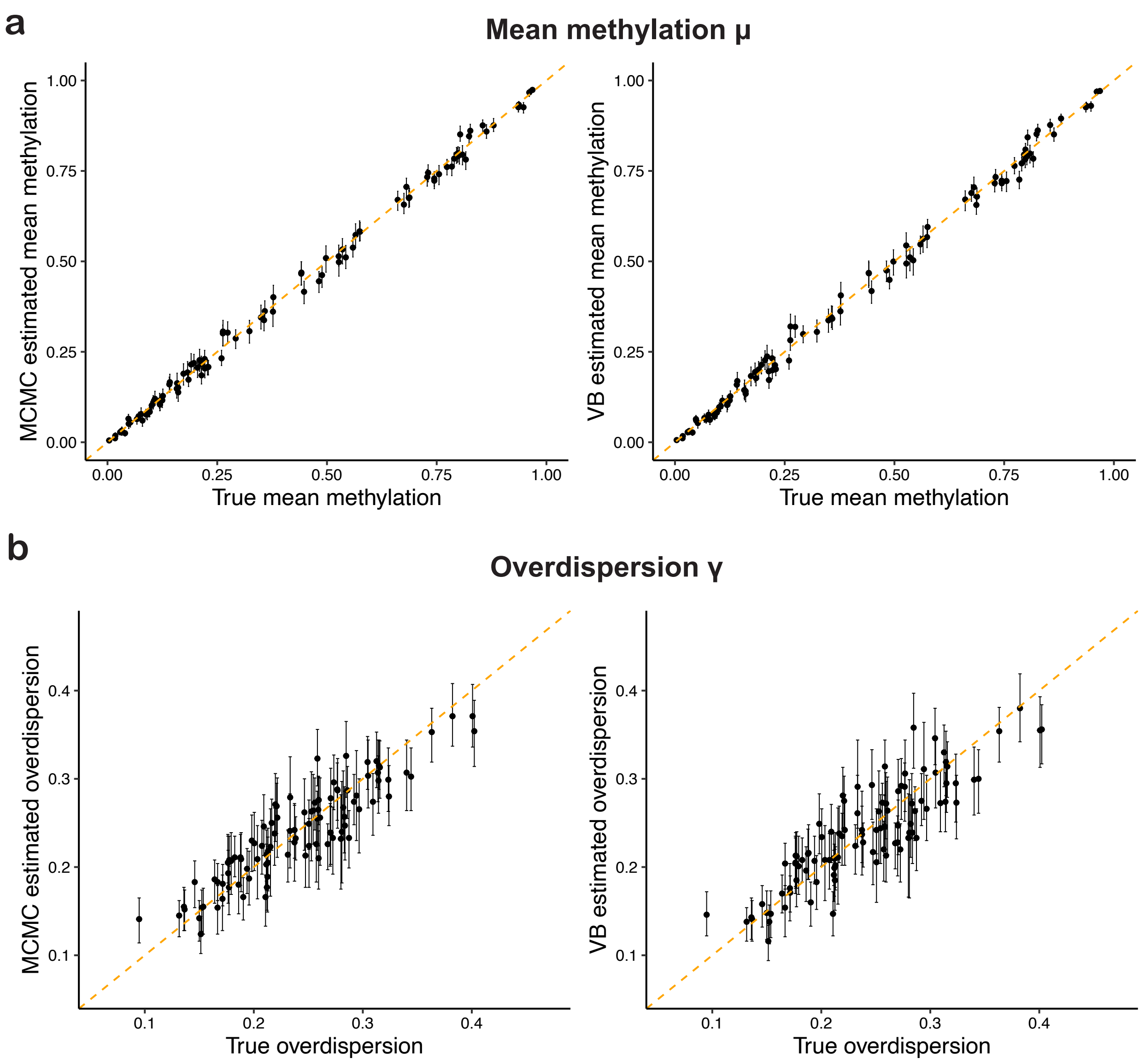


Figure S2: Markov Chain Monte Carlo (MCMC, left panels) and variational Bayes (VB, right panels) inference schemes display comparable estimation performance on simulated data for feature-specific (**a**) mean methylation $\mu_j$ and (**b**) overdispersion $\gamma_j$ parameters. Dashed lines denote perfect agreement between true (x-axis) and estimated (y-axis) parameter values. Each data point represents a different features, dots show posterior medians and vertical lines correspond to 80% high posterior density (HPD) intervals, computed using the `coda` package (Plummer *et al.*, 2006).

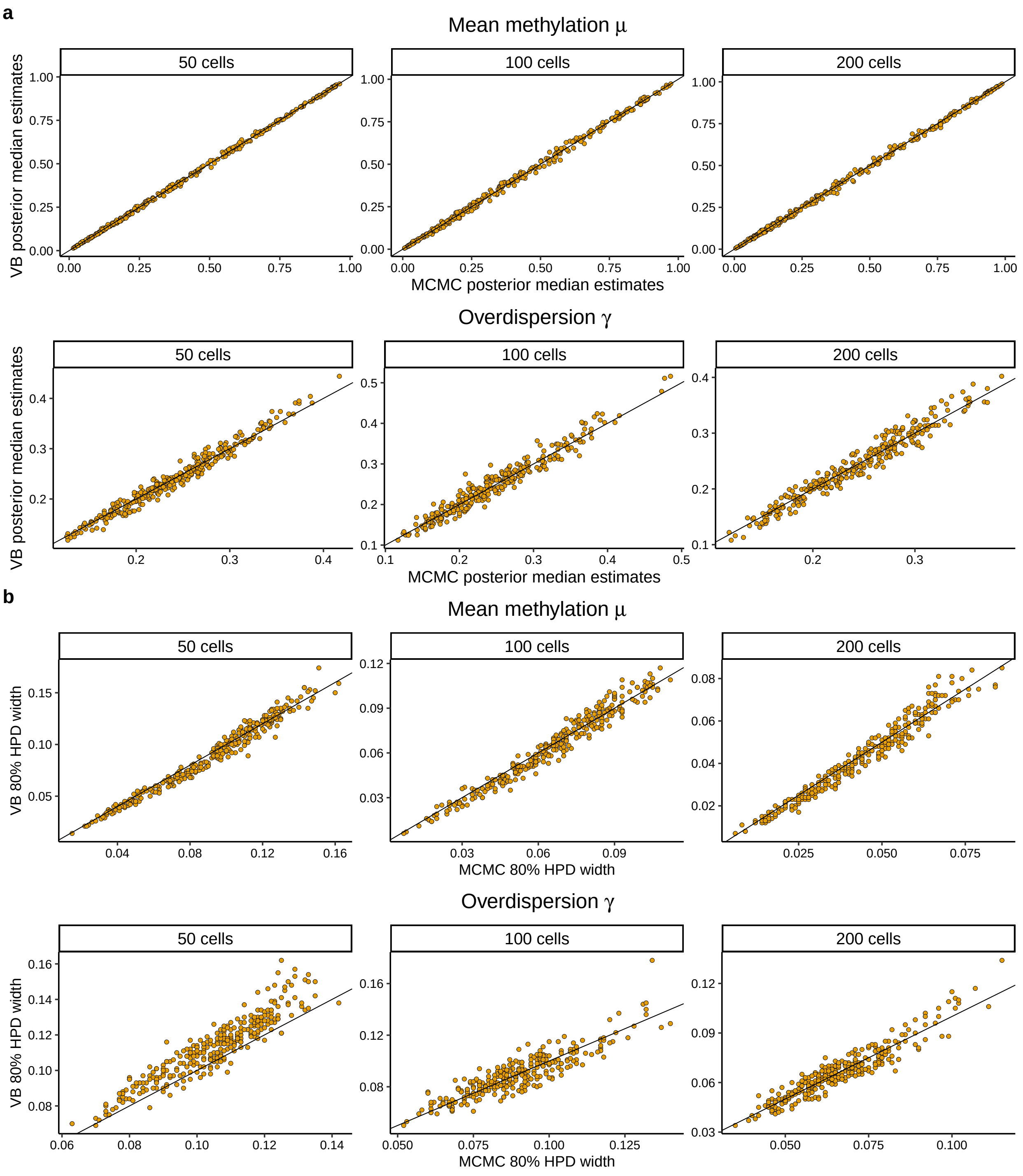


Figure S3: Markov Chain Monte Carlo (MCMC) and variational Bayes (VB) inference schemes infer similar posterior distributions for feature-specific $\mu_j$ and $\gamma_j$ parameters. (**a**) Posterior medians using MCMC (x-axis) and VB (y-axis) for mean methylation (top) and overdispersion (bottom) parameters. (**b**) We compute the width of the 80% high posterior density (HPD) interval using MCMC (x-axis) and VB (y-axis) posterior draws for mean methylation (top) and overdispersion (bottom) parameters. Each data point represents a different feature.

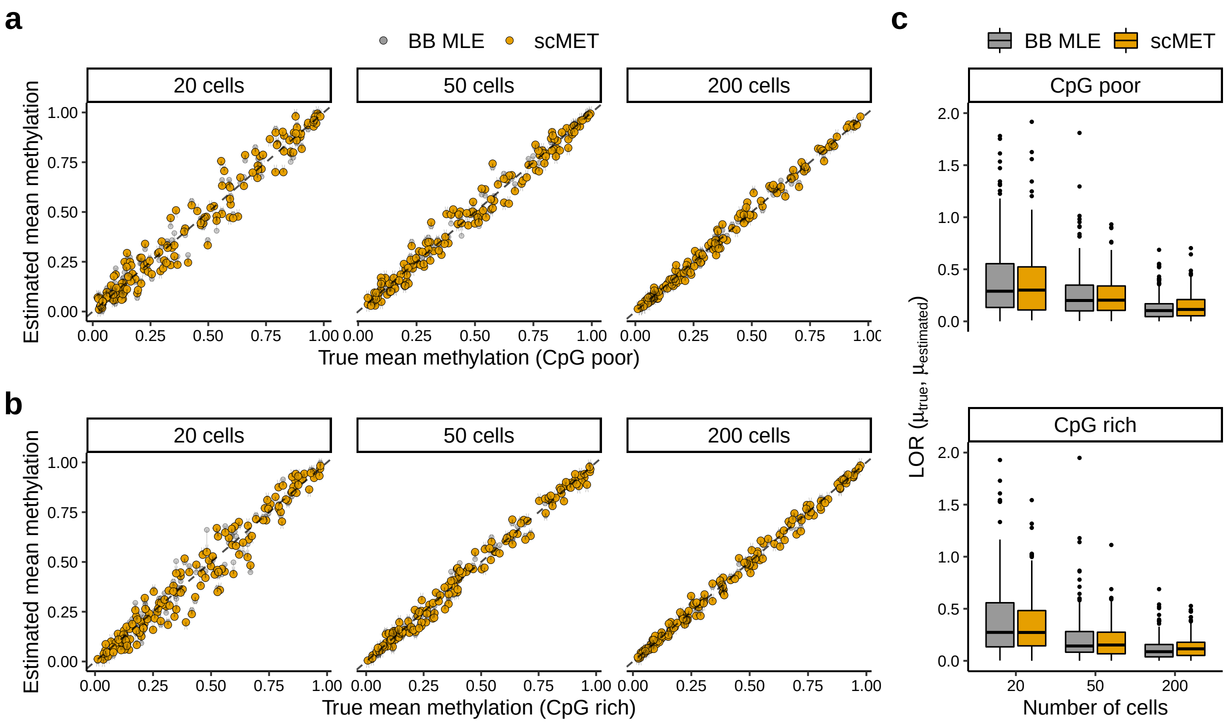


Figure S4: Shrinkage effect on posterior estimates for feature-specific mean methylation parameters $\mu_j$. (**a, b**) Mean methylation estimates per feature for varying number of cells using the beta-binomial MLE (grey) and scMET (yellow) models. For scMET each data point represents the posterior median for the associated parameter. Vertical arrows denote the *shrinkage* that is introduced by scMET with respect to MLE estimates. The dashed line corresponds to perfect agreement between true (x-axis) and estimated (y-axis) mean methylation. As expected, the scMET and BB MLE estimates for $\mu_j$ parameters are comparable. (**a**) CpG poor regions, (**b**) CpG rich regions. (**c**) For the synthetic data in (**a**) and (**b**), we use the absolute log-odds ratio (LOR) difference between true and estimated values as a measure of estimation performance for varying number of cells. The smaller the LOR value the better the estimation performance. Each data point represents a feature.

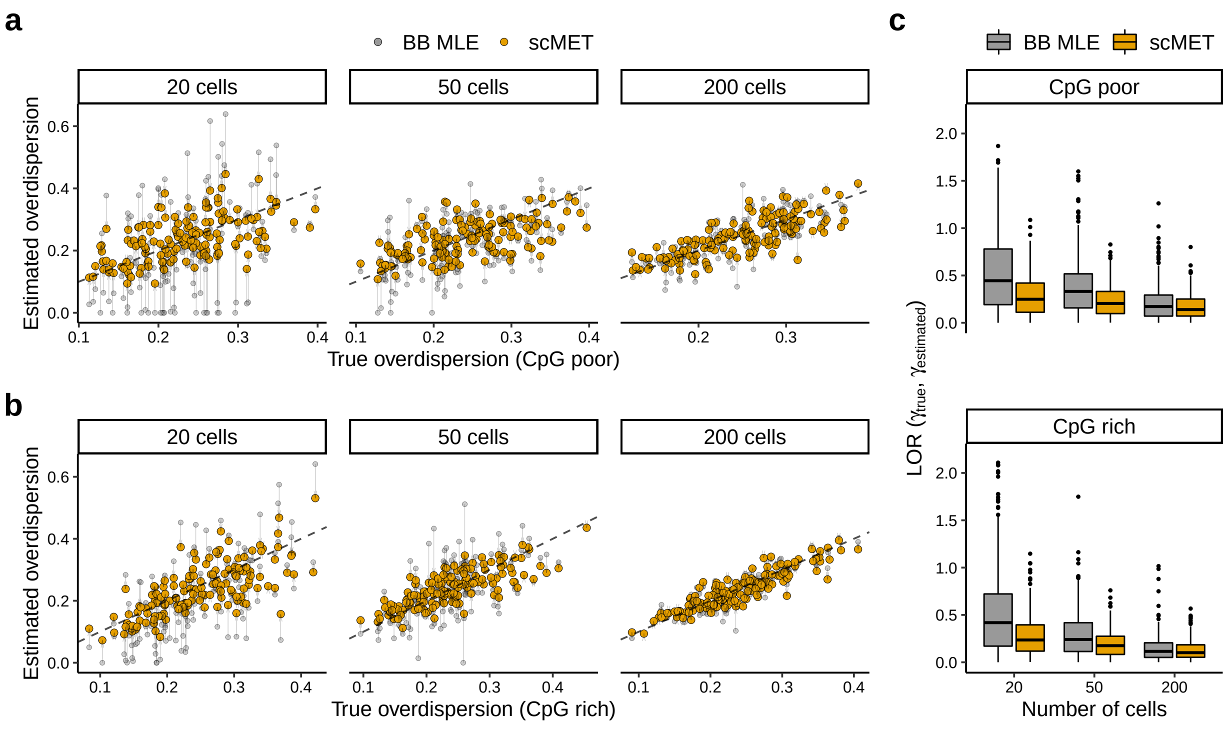


Figure S5: Shrinkage effect on posterior estimates for feature-specific overdispersion parameters $\gamma_j$. (**a, b**) Overdispersion estimates per feature for varying number of cells using the beta-binomial MLE (grey) and scMET (yellow) models. For scMET each point represents the posterior median for the associated parameter. Vertical arrows denote the *shrinkage* that is introduced by scMET with respect to MLE estimates. The dashed line corresponds to perfect agreement between true (x-axis) and estimated (y-axis) overdispersion. (**a**) CpG poor regions. (**b**) CpG rich regions. (**c**) For the synthetic data in (**a**) and (**b**), we use the absolute log-odds ratio (LOR) difference between true and estimated values as a measure of estimation performance for varying number of cells. The smaller the LOR value the better the estimation performance. Each data point represents a feature.

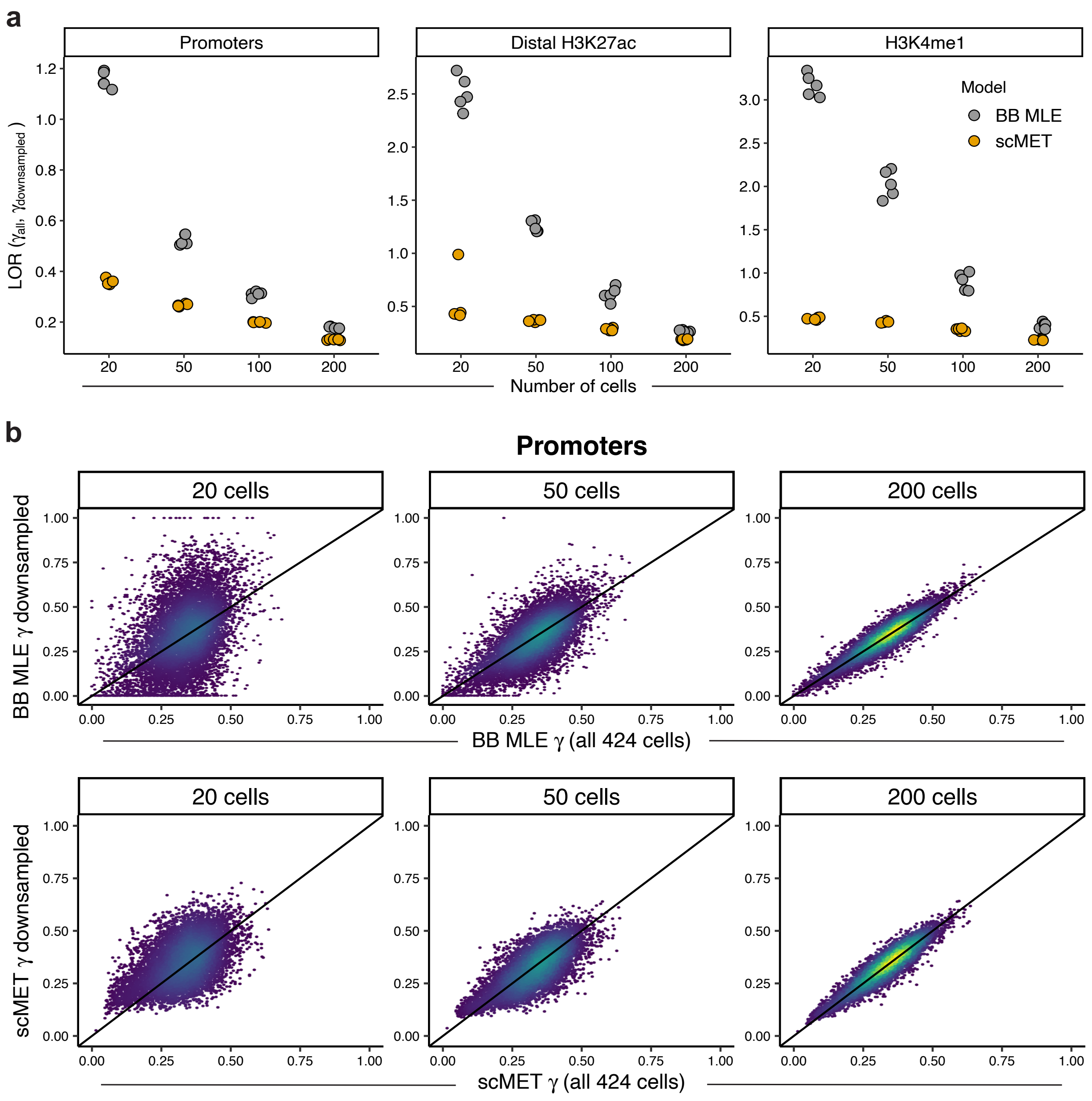


Figure S6: Down-sampling experiments to assess the performance of scMET (posterior medians) and BB MLE point estimates for feature-specific overdispersion parameters $\gamma_j$. In each down-sampling experiment a number of cells are randomly selected amongst the 424 inhibitory neurons characterised by Luo *et al.* (2017). Estimates based on the full dataset are used as pseudo ground truth to assess the stability of inference for decreasing sample size. (**a**) Mean absolute log-odds ratio (LOR) difference as a measure of estimation performance for varying number of cells across the three genomic contexts considered in this study. The smaller the LOR value the better the estimation performance. Each data point represents a different random down-sampling experiment. (**b**) Estimates for feature-specific overdispersion parameters $\gamma_j$ for varying number of cells using BB MLE (top) and scMET (bottom). Each facet compares point estimates of all 424 cells (x-axis) versus a single randomly down-sampled dataset (y-axis). Each data point corresponds to a different feature. The color code within the scatter-plots is used to represent areas with high (green and yellow) and low (blue) concentration of features. Promoter regions are shown as illustrative example; the remaining genomic contexts show a similar pattern (data not shown).

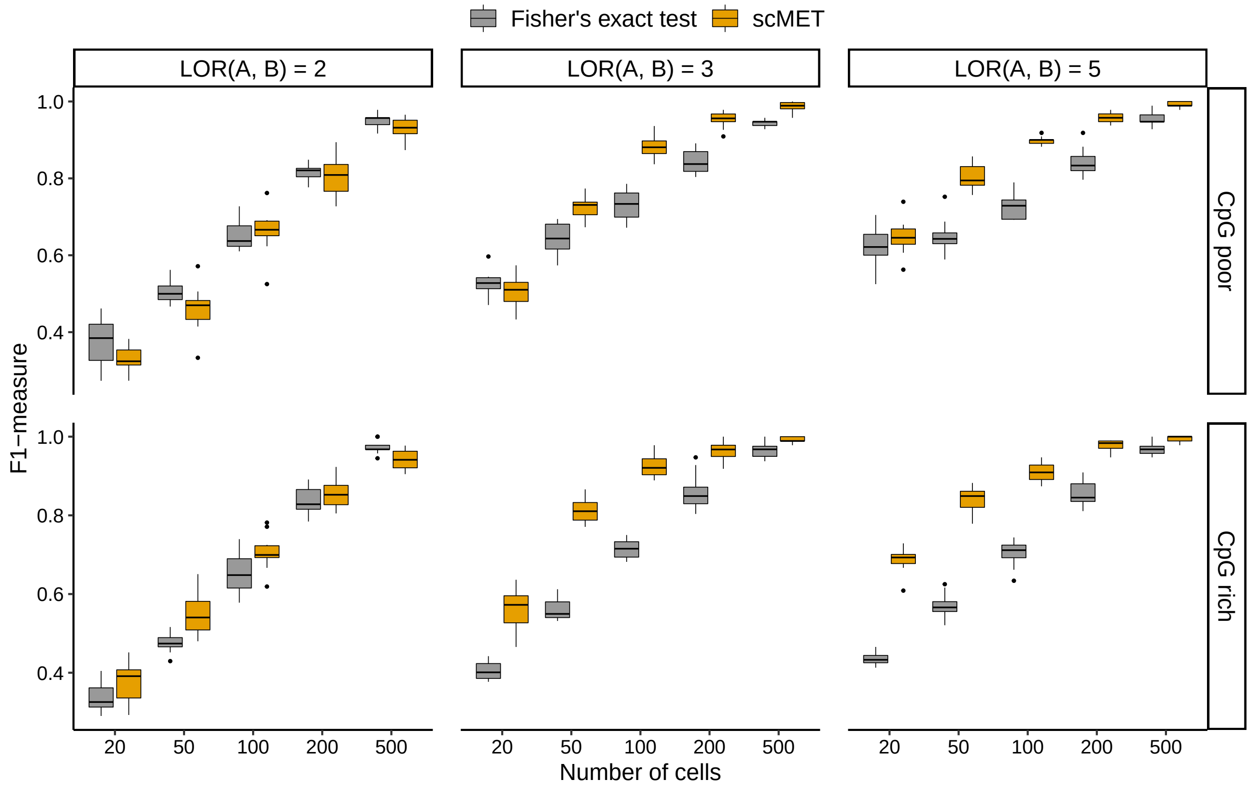


Figure S7: Performance for the differential mean methylation test in terms of F1-measure. We compare hits obtained by scMET (yellow) and Fisher's exact test (grey) in accurately identifying differentially methylated features for varying number of cells (x-axis) and across different settings. Column facets (boxes) correspond to simulating differentially methylated features with different effect sizes, in terms of log-odds ratio (LOR). Row facets correspond to simulated datasets with CpG poor and CpG rich genomic regions (*Methods*). Each data point represents a different synthetic dataset.

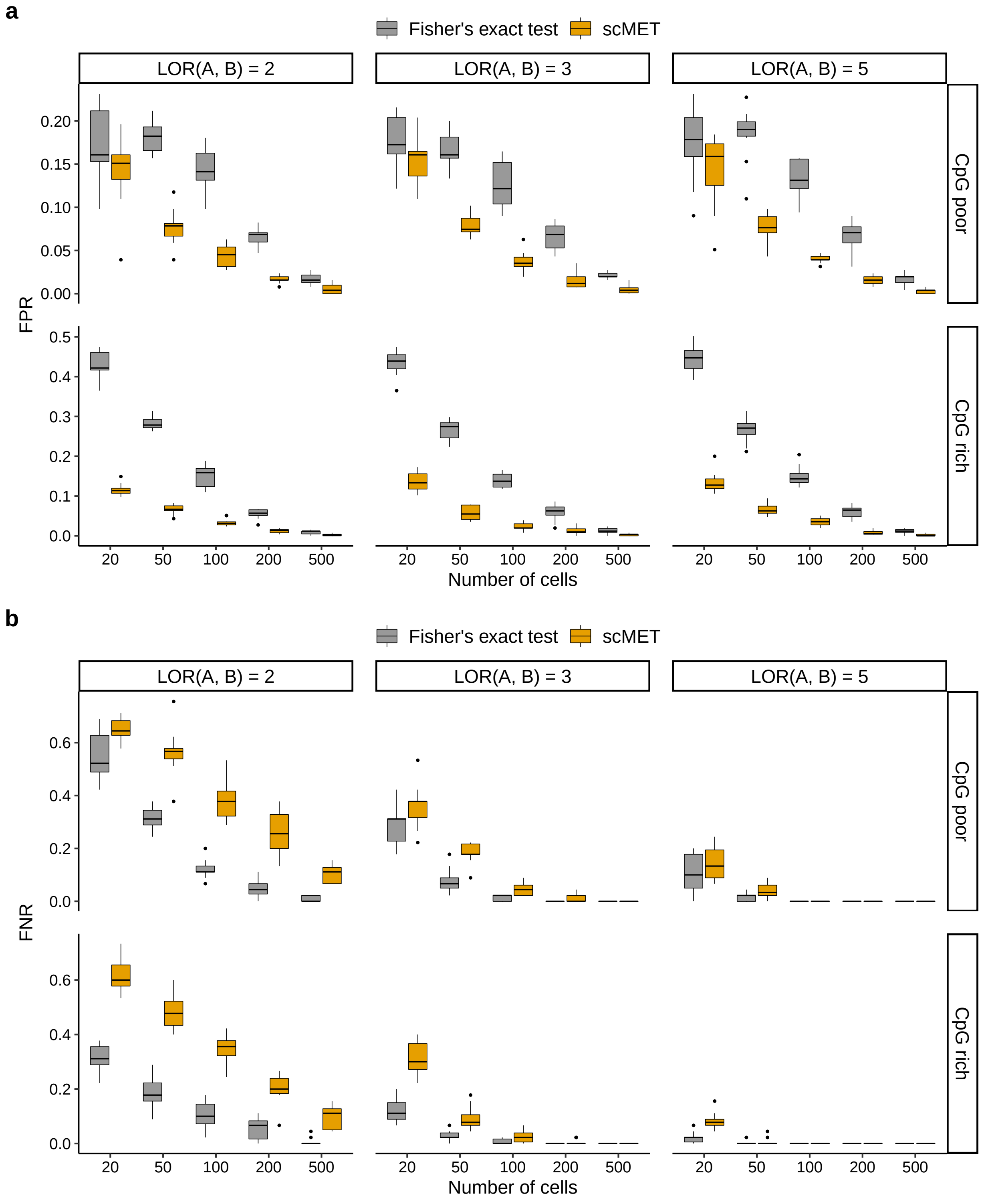


Figure S8: Performance for the differential mean methylation test in terms of false positive rate (FPR, **a**) and false negative rate (FNR, **b**). We compare hits obtained by scMET (yellow) and Fisher's exact test (grey) in accurately identifying differentially methylated features for varying number of cells (x-axis) and across different settings. Column facets (boxes) correspond to simulating differentially methylated features with different effect sizes, in terms of log-odds ratio (LOR). Row facets correspond to simulated datasets with CpG poor and CpG rich genomic regions (*Methods*). Each data point represents a different synthetic experiment.

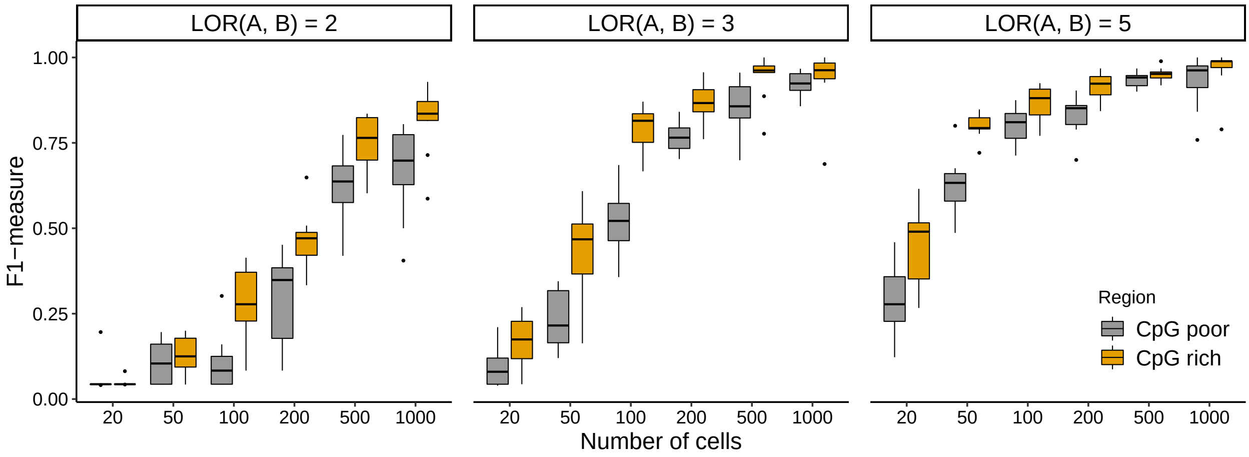


Figure S9: Performance for the differential variability test in terms of F1-measure (y-axis) for varying number of cells (x-axis). Colours correspond to simulated datasets with CpG poor (grey) and CpG rich (yellow) genomic regions (*Methods*). Column facets (boxes) correspond to simulating differentially variable features with different effect sizes, in terms of log-odds ratio (LOR). Each data point represents a different synthetic experiment.

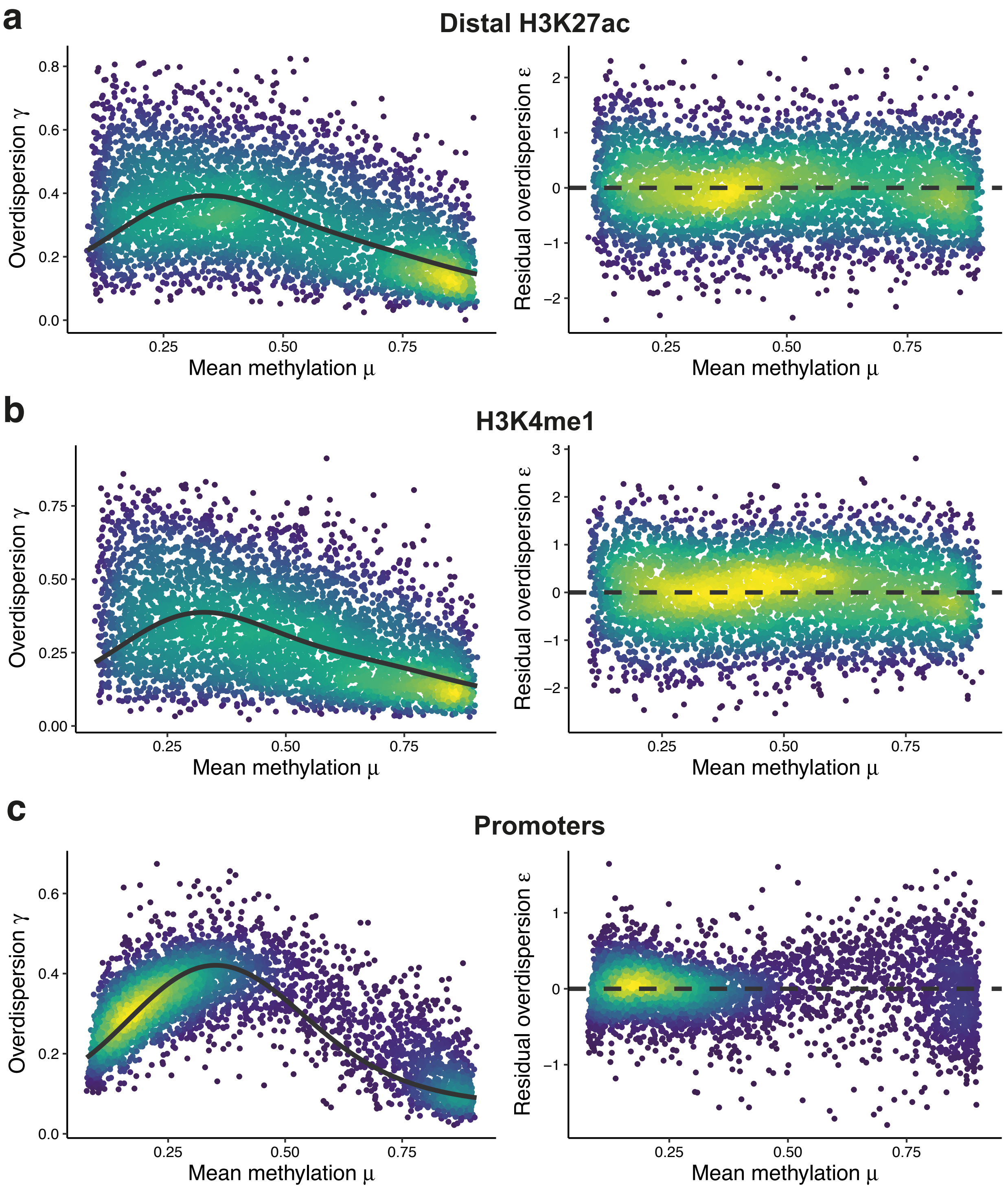


Figure S10: Mean-overdispersion relationship across genomic contexts for the Luo *et al.* (2017) dataset. Each horizontal subpanel corresponds to a different genomic context. Left panels show posterior medians for feature-specific mean and overdispersion parameters. Black line represents the estimated regression trend from the GLM component of scMET (see Fig. 1a). Right panels show posterior medians for feature-specific methylation parameters $\mu_j$ (x-axis) versus residual overdispersion parameters $\epsilon_j$ (y-axis). Each data point corresponds to a different feature. The color code within the scatter-plots is used to represent areas with high (green and yellow) and low (blue) concentration of features.

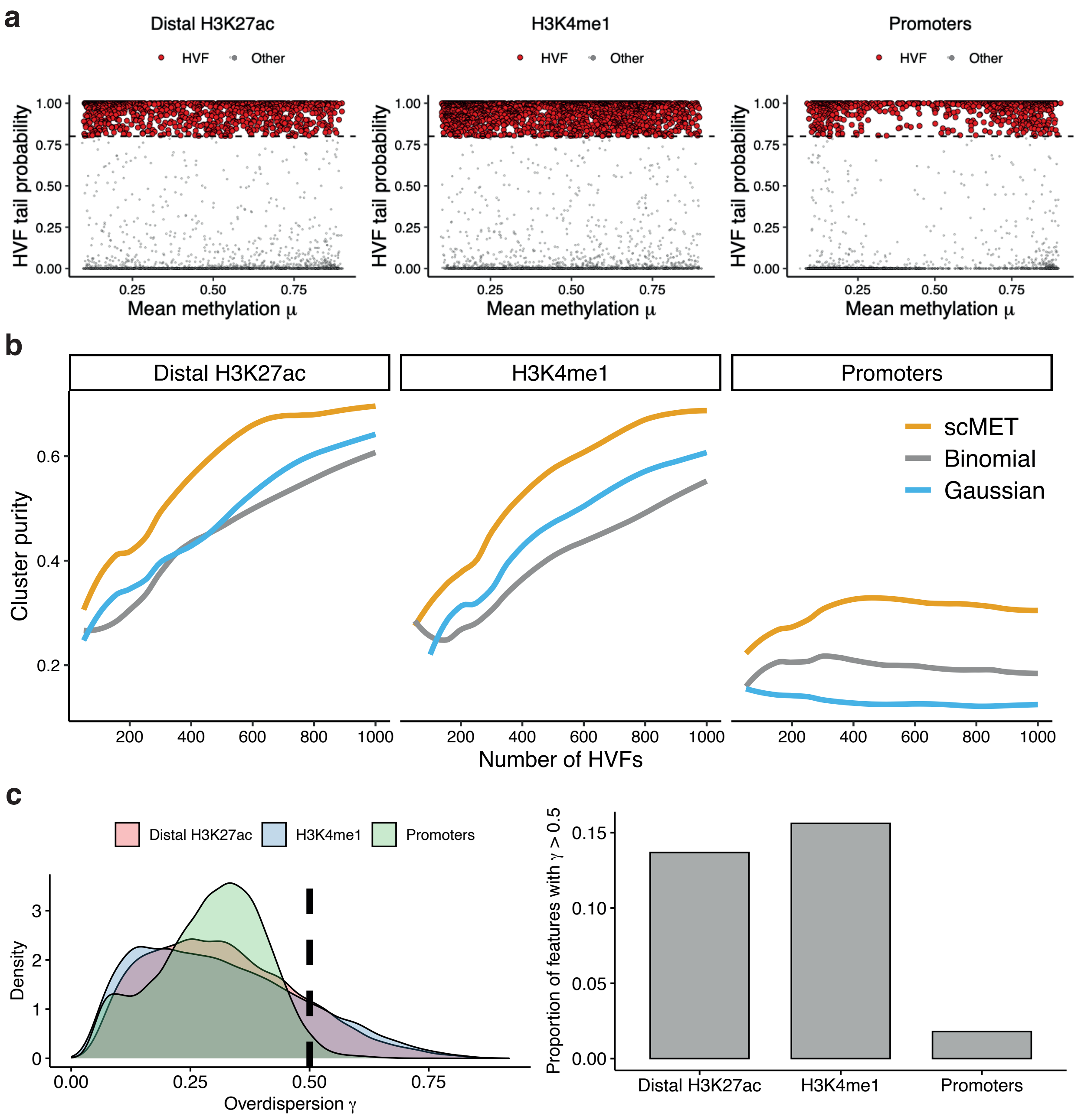


Figure S11: Identifying highly variable features (HVFs). (**a**) scMET tail posterior probabilities denoting the evidence of a feature being called as HVF. Each panel corresponds to a different genomic context for the Luo *et al.* (2017) dataset. Red points correspond to features that are labelled as HVF. The horizontal dashed line corresponds to the posterior evidence threshold $\alpha_H$ set to match a desired EFDR = 10% (*Methods*). (**b**) Clustering performance, in terms of cluster purity (see Section S2.3), for varying number of selected HVFs. HVF selection was based on scMET (yellow), binomial variance (grey) and Gaussian variance (blue). A finite grid of HVFs was used for cluster purity evaluation and non-parametric regression was used to obtain a smoothed interpolation across all values (*Methods*) (**c**) Distribution of posterior estimates for feature-specific overdispersion parameters $\gamma_j$ across different genomic contexts (left). Percentage of features with $\gamma_j > 0.5$ per genomic context (right).

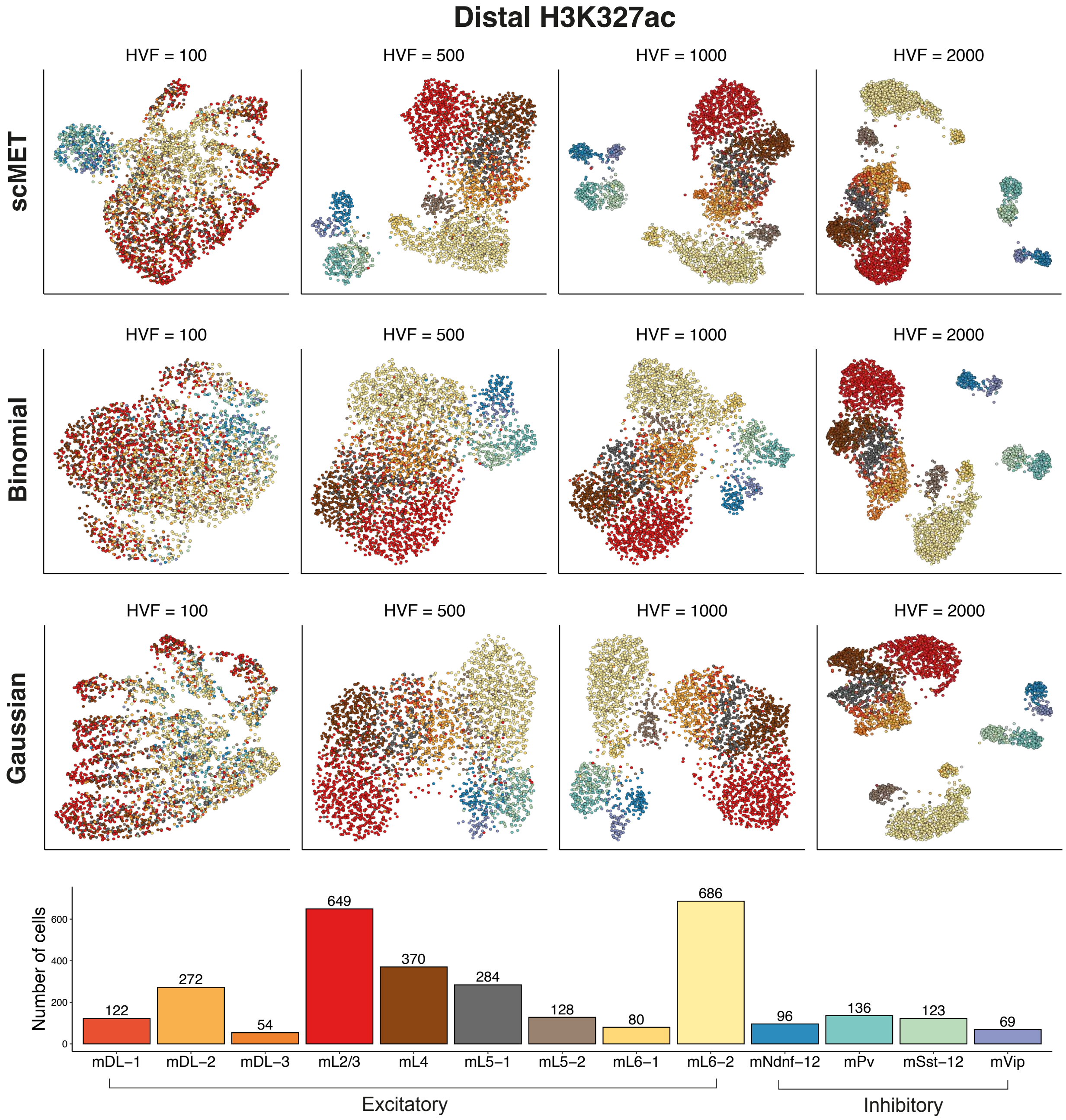


Figure S12: UMAP representations for varying number of HVFs when applied to distal H3K27ac features on the Luo *et al.* (2017) dataset. Each row corresponds to a different method for HVF selection (scMET, Binomial and Gaussian). Each column corresponds to different number of HVFs. Each data point corresponds to a different cell. Points are coloured according to cell type assignment in the Luo *et al.* (2017) study as shown in the bottom panel.

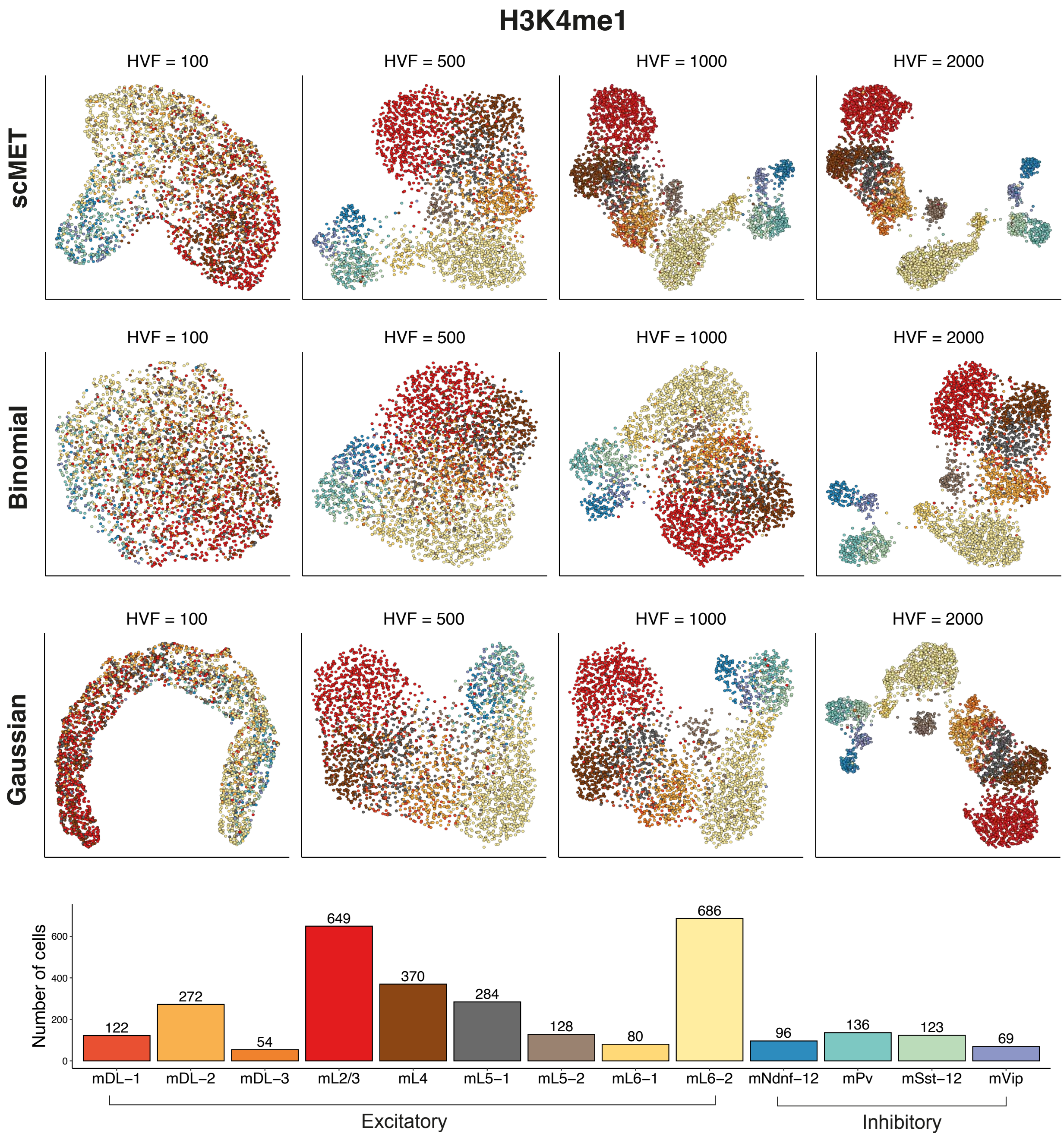


Figure S13: UMAP representations for varying number of HVFs when applied to H3K4me1 features on the Luo *et al.* (2017) dataset. Each row corresponds to a different method for HVF selection (scMET, Binomial and Gaussian). Each column corresponds to different number of HVFs. Each data point corresponds to a different cell. Points are coloured according to cell type assignment in the Luo *et al.* (2017) study as shown in the bottom panel.

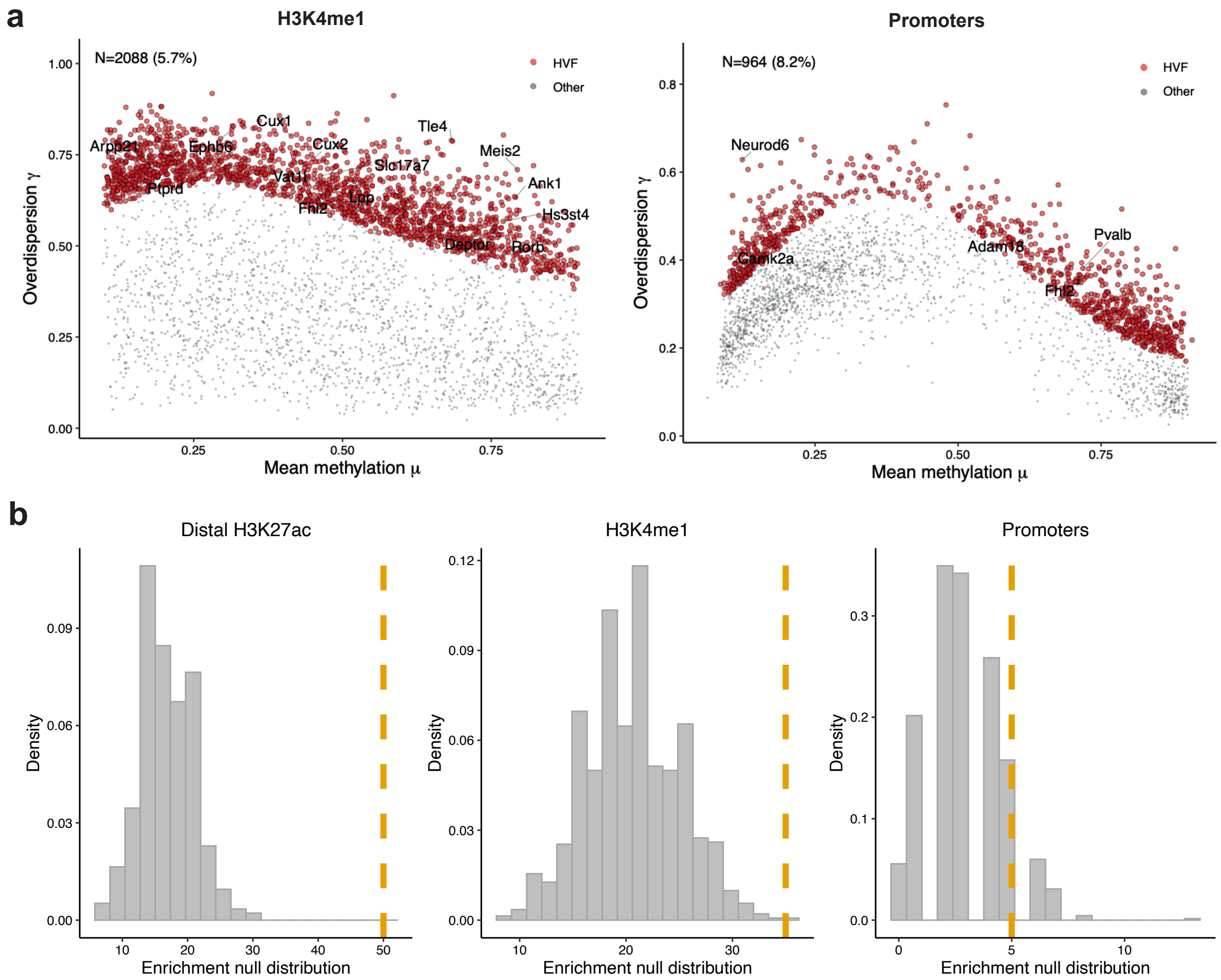


Figure S14: Identifying highly variable features (HVFs) on the Luo *et al.* (2017) dataset. (**a**) Mean-overdispersion plots for H3K4me1 and promoter genomic contexts. Red points correspond to features being called as HVF (EFDR = 10% and percentile threshold $\delta_E = 90\%$). To ease interpretation each element is linked to its nearest gene. (**b**) Histogram showing enrichment distribution under the null model. The null model is based on randomly labelling features as HVFs and then counting how many of those are overlapping with neuron marker genes identified by Luo *et al.* (2017), see Supplementary Table S1. To obtain a null distribution we repeated this process 1,000 times. The dashed vertical yellow line, shows the enrichment of HVFs using the scMET model.

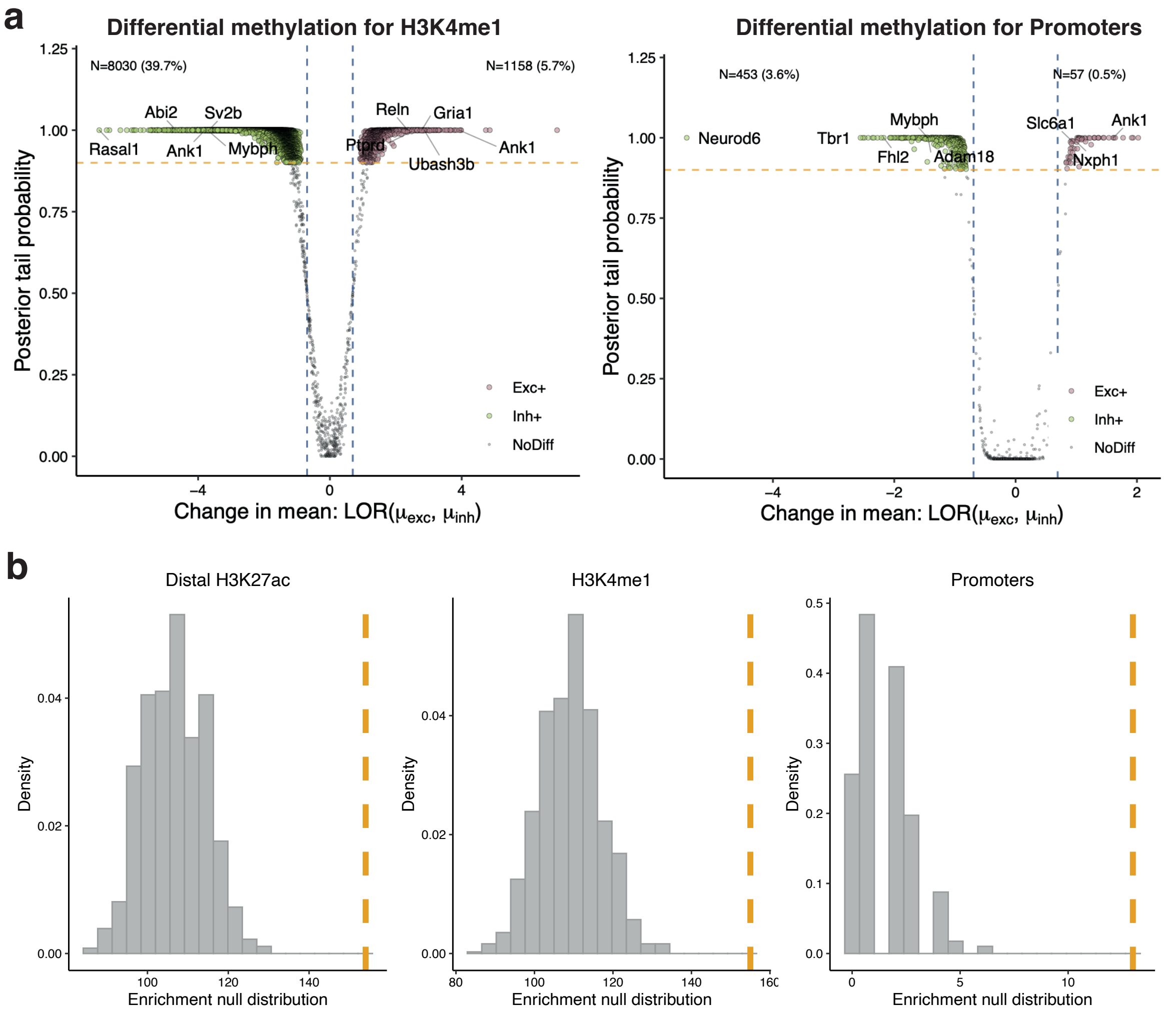


Figure S15: Identifying differentially methylated (DM) features across excitatory and inhibitory neurons on the Luo *et al.* (2017) dataset. (**a**) Volcano plots of DM features for H3K4me1 and promoter genomic contexts. (**b**) Histogram showing enrichment distribution under the null model. The null model is based on randomly labelling features as DM and then counting how many of those are overlapping with neuron marker genes identified in Luo *et al.* (2017), see Supplementary Table S1. To obtain a null distribution we repeated this process 1,000 times. The dashed vertical yellow line, shows the enrichment of DM features using the scMET model.

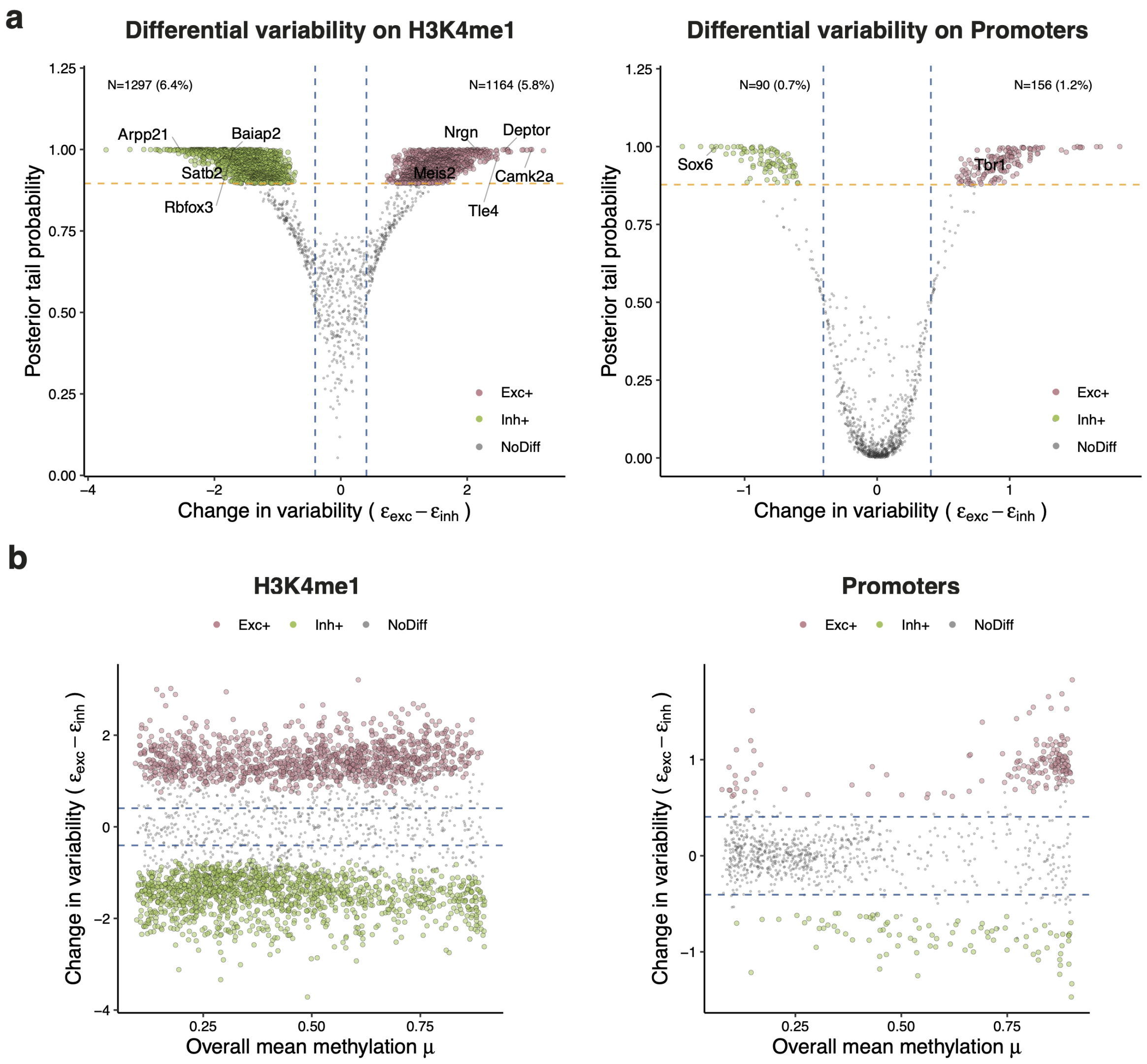


Figure S16: Identifying differentially variable (DV) features across excitatory and inhibitory neurons on the Luo *et al.* (2017) dataset. (**a**) Volcano plots of DV features for H3K4me1 and promoter genomic contexts. (**b**) For each feature, the overall mean methylation across all cells is plotted against the change in residual overdispersion between excitatory and inhibitory neurons. Features with statistically significant changes in variability are coloured according to their regulation. Red-like and green-like colours denote features with higher variability in excitatory and inhibitory neurons, respectively.

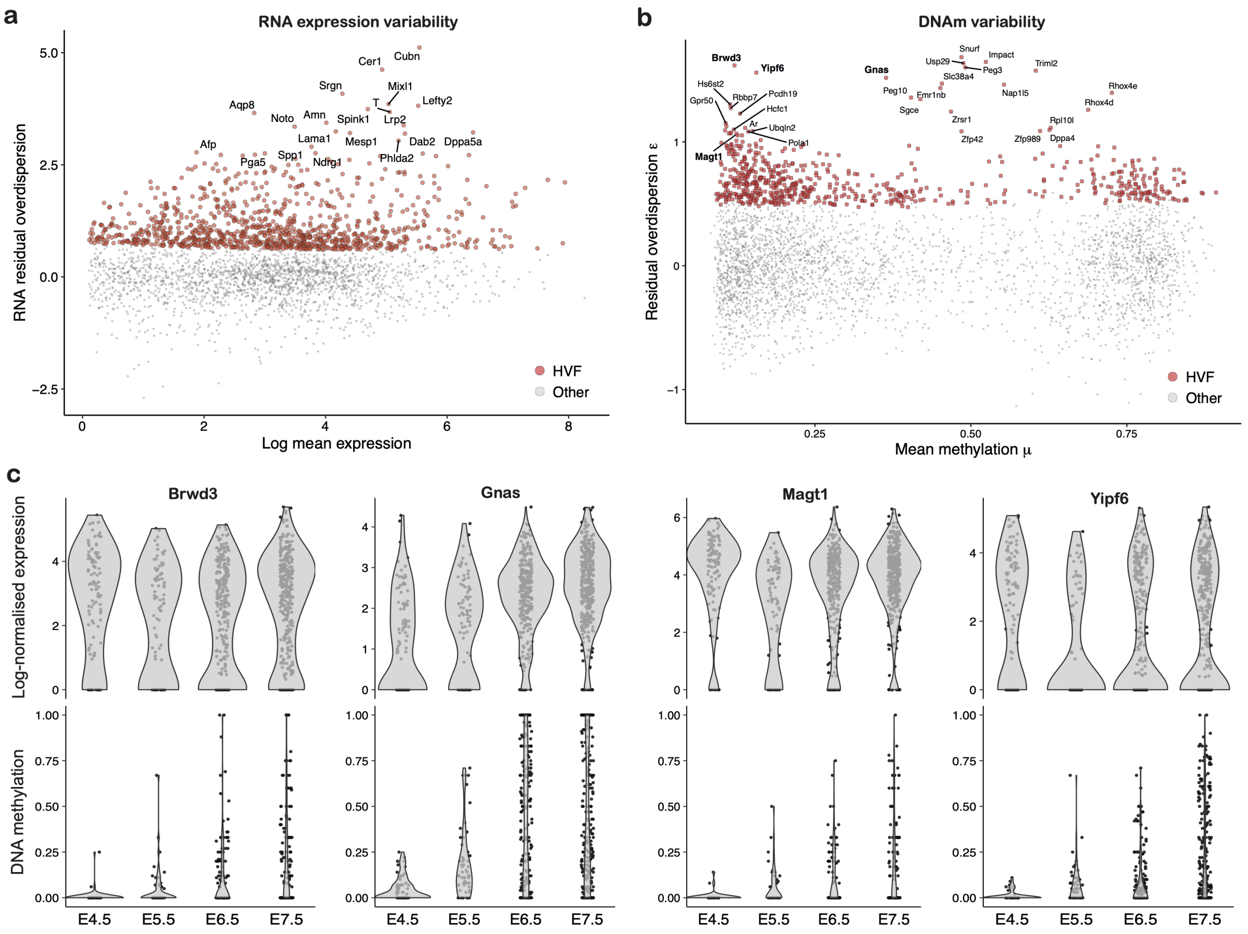


Figure S17: Exploring the relationship between transcriptional and DNAm variability on the multi-omics scNMT-seq gastrulation dataset (Argelaguet *et al.*, 2019). Mean (x-axis) versus residual overdispersion estimates (y-axis) for (**a**) RNA expression and (**b**) promoter DNA methylation modalities for the scNMT-seq gastrulation dataset. (**c**) Representative examples of the DNAm and RNA expression patterns for genes with high DNA methylation heterogeneity and low RNA expression heterogeneity. Y-axis shows BASiCS log-normalised gene expression (in a $\log(x + 1)$ scale) (top) and promoter DNAm rate (bottom). Cells are stratified by embryonic stage (x-axis).

### S2 Supplementary notes

#### S2.1 scMET prior specification

To complete the scMET model we introduce the following priors for the remaining parameters,

$$\mathbf{w}_\mu \sim \text{MVN}(\mathbf{m}_{w\mu}, s_{w\mu}\mathbf{I}), \quad \mathbf{w}_\gamma \sim \text{MVN}(\mathbf{m}_{w\gamma}, s_{w\gamma}\mathbf{I}), \quad s_\gamma \sim \text{IG}(\alpha_{s\gamma}, \beta_{s\gamma}),$$

where MVN$(\cdot)$ denotes the multivariate-normal distribution, IG$(\cdot)$ the inverse-gamma distribution, and $\mathbf{I}$ the identity matrix with appropriate dimensions. The joint distribution over the observed and latent variables for the scMET model is given by,

$$\begin{aligned} p(\mathbf{Y}, \boldsymbol{\mu}, \boldsymbol{\gamma}, \mathbf{w}_\mu, s_\mu, \mathbf{w}_\gamma, s_\gamma \,|\, \mathbf{X}) = p(\mathbf{Y} \,|\, \boldsymbol{\mu}, \boldsymbol{\gamma})\; p(\boldsymbol{\mu} \,|\, \mathbf{w}_\mu, s_\mu, \mathbf{X}) \\ p(\boldsymbol{\gamma} \,|\, \boldsymbol{\mu}, \mathbf{w}_\gamma, s_\gamma)\; p(\mathbf{w}_\mu)\; p(\mathbf{w}_\gamma)\; p(s_\gamma), \end{aligned} \tag{1}$$

where the factorisation corresponds to the probabilistic graphical model provided in Fig. 1a and Supplementary Fig. S18.

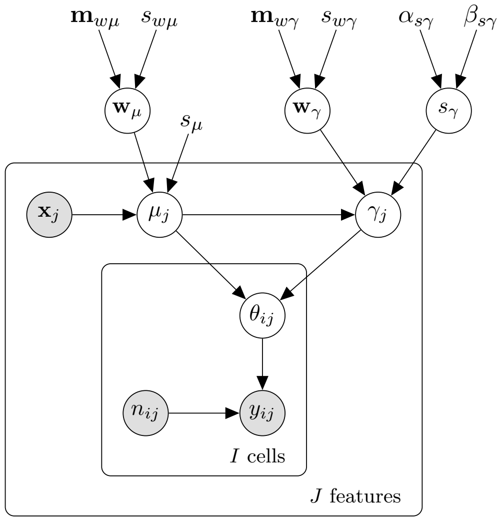


$$\begin{aligned} s_\gamma &\sim \text{IG}(\alpha_{s\gamma}, \beta_{s\gamma}) \\ \mathbf{w}_\gamma &\sim \mathcal{N}(\mathbf{m}_{w\gamma}, s_{w\gamma}) \\ \mathbf{w}_\mu &\sim \mathcal{N}(\mathbf{m}_{w\mu}, s_{w\mu}) \\ \gamma_j | \mu_j, \mathbf{w}_\gamma, s_\gamma &\sim \text{Logit}\mathcal{N}\big(f_\gamma(\mu_j; \mathbf{w}_\gamma), s_\gamma\big) \\ \mu_j | \mathbf{x}_j, \mathbf{w}_\mu, s_\mu &\sim \text{Logit}\mathcal{N}\big(f_\mu(\mathbf{x}_j; \mathbf{w}_\mu), s_\mu\big) \\ \theta_{ij} | \mu_j, \gamma_j &\sim \text{Beta}(\mu_j, \gamma_j) \\ y_{ij} | \theta_{ij}, n_{ij} &\sim \text{Binomial}(n_{ij}, \theta_{ij}) \\ f_\mu(\mathbf{x}_j; \mathbf{w}_\mu) &= \mathbf{w}_\mu^\top \mathbf{x}_j \\ f_\gamma(\mu_j; \mathbf{w}_\gamma) &= w_{\gamma 1} + \sum_{l=2}^{L} w_{\gamma l} g_l(\mu_j) \end{aligned}$$

Figure S18: Probabilistic graphical representation of the scMET model.

##### S2.1.1 Choice of hyper-parameters

For all experiments performed in this study the following hyper-parameters were fixed *a priori* to the following default values:

$$s_\mu = 1.5, \quad s_{w\mu} = 2, \quad s_{w\gamma} = 2, \quad \alpha_{s\gamma} = 2, \quad \beta_{s\gamma} = 3$$

For the $\mathbf{m}_{w\mu}$ and $\mathbf{m}_{w\gamma}$ hyper-parameters we employed an empirical Bayes approach (Gelman *et al.*, 2013) to set default values based on the data. To do so, we first obtained MLE estimates $\hat{\mu}_j$ and $\hat{\gamma}_j$ using the VGAM package. Then, $\mathbf{m}_{w\mu}$ values were set as the coefficients of the following linear regression model: logit$(\hat{\mu}_j) = \mathbf{m}_{w\mu}^\top \mathbf{x}_j + \epsilon_j$. For all analyses we assumed the feature-specific covariates were $\mathbf{x}_j = (1, C_j)$, where $C_j$ denotes the CpG density. The $\mathbf{m}_{w\gamma}$ values were set as the coefficients of the following basis function regression model:

$$\text{logit}(\hat{\gamma}_j) = m_{w\gamma 1} + \sum_{l=2}^{L} m_{w\gamma l} g_l(\mu_j) + \epsilon_j \tag{2}$$

where $g_l(\cdot)$ are radial basis functions as defined in Kapourani and Sanguinetti (2016). The total number of basis functions $L$ was fixed to 4 for all analyses; a choice flexible enough to capture the mean-variance relationship present in the data analysed in this study.

#### S2.2 EFDR calibration

The posterior evidence thresholds $\alpha$ quantify the uncertainty associated with the differential test or HVF analysis and can be fixed *a priori*. Otherwise, we can choose optimal thresholds to control the expected false discover rate (EFDR, Newton *et al.*, 2004) given by,

$$\text{EFDR}(\alpha) = \frac{\sum_{j=1}^{J} \big(1 - \pi_j(\psi)\big)\mathbb{I}\big(\pi_j(\psi) > \alpha\big)}{\sum_{j=1}^{J} \mathbb{I}\big(\pi_j(\psi) > \alpha\big)}, \tag{3}$$

where $\mathbb{I}(s) = 1$ if $s$ is true, otherwise 0, $\alpha$ denotes the posterior evidence threshold and $\psi$ the posterior probability. Unless otherwise stated, for DM and DV analysis we set EFDR = 5% and for HVF analysis we set EFDR = 10%. As discussed in Vallejos *et al.* (2016), the usability of this calibration rule relies on the existence of features under both the null and alternative hypothesis (i.e. with and without changes in methylation patterns). As a default, if EFDR calibration is not achieved, we set $\alpha = 0.9$.

#### S2.3 Miscellaneous

**Logit.** If $p \in (0, 1)$ is a probability, the logit or log-odds function is defined as the logarithm of the odds,

$$\text{logit}(p) = \log\left(\frac{p}{1-p}\right). \tag{4}$$

**Log-odds ratio.** The logarithm of the odds ratio (LOR) is equal to the difference between the logits of two probabilities, that is:

$$\text{LOR}(p_A, p_B) = \log\left(\frac{p_A/(1-p_A)}{p_B/(1-p_B)}\right) = \text{logit}(p_A) - \text{logit}(p_B). \tag{5}$$

**F1-measure.** The F1-measure or F1-score is the harmonic mean of precision and recall:

$$\text{F1-measure} = 2 \cdot \frac{\text{precision} \cdot \text{recall}}{\text{precision} + \text{recall}}. \tag{6}$$

**Cluster purity.** Purity is a measure of the extent to which clusters contain a single class. Given some set of clusters M and some set of classes D, both partitioning N data points, purity can be defined as:

$$\frac{1}{N} \sum_{m \in M} \max_{d \in D} |m \cap D|. \tag{7}$$
