## Supplementary Tables for "scMET: Bayesian modelling of DNA methylation heterogeneity at single-cell resolution": README.txt

Supplementary Tables for scMET: Bayesian modelling of DNA methylation heterogeneity at single cell resolution.

Table S1: Neuronal marker genes used in the Luo et al. (2017) study to characterise the different neuronal sub-populations. Use for enrichment analysis.
Table S2: Sample metadata information for the mouse frontal cortex dataset by Luo et al. (2017).
Table S3: Download information on the two ChIP-seq datasets (H3K27ac and H3K4me1) that were profiled in adult (8 weeks) mouse cortex and are available as part of the ENCODE project.
Table S4: Sample metadata information for the scNMT-seq gastrulation dataset by Argelaguet et al. (2019).
